# Transcriptional programs define solid and ascitic dissemination in ovarian cancer

**DOI:** 10.1101/2025.08.26.672372

**Authors:** Kaiyang Zhang, Liangru Fei, Daria Afenteva, Essi Kahelin, Giovanni Marchi, Yilin Li, Jaana Oikkonen, Sakari Hietanen, Johanna Hynninen, Biswajyoti Sahu, Sampsa Hautaniemi, Anni Virtanen

## Abstract

Metastatic cancer cells must survive in distinct anatomical environments, yet how cancer-cell plasticity is organized across these environments remains poorly understood. Here, we integrated RNA and whole-genome sequencing of 600 multiregion and longitudinal specimens from 284 patients with ovarian high-grade serous carcinoma (HGSC), complemented by DNA methylation, single-nucleus multiome profiling and quantitative histomorphology. Among 25 recurrent cancer-cell transcriptional programs, Mes-ECM was associated with solid omental and peritoneal metastases, whereas Mechano-EMP was associated with ascites. Importantly, these compartment-associated transcriptional programs were already present in treatment-naïve tubo-ovarian tumors, where their activity was linked to transcriptional compatibility with solid metastatic sites or ascites. Despite their distinct anatomical associations, both states shared a TEAD–AP-1/YAP–TAZ mechanotransductive signature but diverged in their regulatory and microenvironmental relationships. Mes-ECM was linked to extracellular-matrix remodeling, outward stromal signaling and cancer-proximal stromal expansion, whereas Mechano-EMP integrated hybrid epithelial–mesenchymal and stress–inflammatory programs with SDC4-centered inputs from stromal, vascular and myeloid cells. Both programs were associated with altered collagen organization, while chemotherapy-associated remodeling and prognostic associations varied by anatomical site. Our findings reveal a modular organization of metastatic cancer-cell plasticity, linking shared regulatory circuitry to distinct anatomical environments and tumor ecosystems.

## INTRODUCTION

Metastasis is the leading cause of cancer death, yet how cancer cells adapt as they disseminate and colonize distinct anatomical sites remains incompletely understood ^1–3^. Peritoneal dissemination, a hallmark of advanced disease in several intra-abdominal malignancies, provides a particularly informative setting to investigate metastatic adaptation, as cancer cells must survive in peritoneal fluid and establish metastases in solid tissues ^4^. Ovarian high-grade serous carcinoma (HGSC) exemplifies this process, spreading predominantly through the transcoelomic route, with tubo-ovarian tumors, solid omental and peritoneal metastases, and malignant ascites frequently coexisting within the same patient ^5,6^. These distinct anatomical compartments impose different biological requirements on cancer cells, raising the question of whether intraperitoneal dissemination involves a common metastatic cell state or distinct plasticity programs associated with survival in ascites and colonization of solid tissues.

Multiregion genomic studies have established a complex evolutionary history of HGSC, revealing non-random patterns of clonal dissemination and distinct evolutionary states associated with molecular and clinical phenotypes ^7–9^. Across other cancers, untreated metastases often harbor few metastasis-private clonal mutations, while functional driver mutations are largely conserved among a patient’s metastatic lesions ^10,11^. Although these studies have elucidated the clonal ancestry and evolutionary trajectories of metastatic tumors, they do not establish the cancer-cell states associated with adaptation to distinct anatomical environments. Growing evidence across cancers indicates that metastatic progression involves developmental, epithelial–mesenchymal, and stress-adaptive transcriptional programs shaped by interactions with the tumor microenvironment (TME) ^1,2^. In HGSC, recent studies have identified anatomical site-associated differences in immune and cancer-cell phenotypes ^12,13^. However, how recurrent cancer-cell plasticity programs relate to dissemination across solid metastatic tissues and ascites, and what regulatory and microenvironmental features distinguish these programs, remains poorly understood.

Herein, we investigated how cancer-cell plasticity relates to intraperitoneal dissemination in HGSC using 600 multiregion and longitudinal specimens from 284 patients. We identified recurrent cancer-cell transcriptional programs and uncovered distinct forms of plasticity associated with solid metastases and ascites, with corresponding transcriptional features already present in tubo-ovarian tumors. These states shared a core mechanotransductive regulatory program but differed in their regulatory circuitry, interactions with the TME, and associated tissue architecture. Together, our findings reveal how distinct configurations of cancer-cell plasticity are linked to the different biological demands of solid-tissue colonization and persistence in ascites.

## RESULTS

### Multimodal profiling dissects the HGSC transcriptional landscape

To characterize the transcriptional landscape of HGSC across anatomical sites and treatment phases, we analyzed 600 tumor samples from 284 patients enrolled in the observational, prospective, longitudinal, multiregion DECIDER clinical trial (ClinicalTrials.gov: NCT04846933). Most patients had advanced-stage disease (70.1% FIGO stage III and 26.8% stage IV). Primary debulking surgery followed by platinum-based chemotherapy was performed in 54.6% of patients, whereas 45.4% received neoadjuvant chemotherapy (NACT). Median overall survival was 38 months (Table 1).

**Table 1.** Summary of the demographics and clinical features of patients in the study.

| **Feature** | **All patients (284)** |
|---|---|
| Age at diagnosis |  |
| Median | 69 |
| IQR | 64-74 |
| FIGO 2014 Stage, n (%) |  |
| I | 2 (0.7%) |
| II | 7 (2.5%) |
| III | 199 (70.1%) |
| IV | 76 (26.8%) |
| Treatment strategy, n (%) |  |
| PDS | 155 (54.6%) |
| NACT | 129 (45.4%) |
| Residual disease, n (%) |  |
| 0 | 110 (38.7%) |
| 1 to 10mm | 106 (37.3%) |
| > 10mm | 41 (14.4%) |
| No debulking surgery | 27 (9.5%) |
| Pretreatment SBS3 status, n (%) |  |
| Negative | 133 (46.8%) |
| Positive | 107 (37.7%) |
| Not available | 44 (15.5%) |
| Maintenance therapy after 1st line chemotherapy, n (%) |  |
| No | 123 (43.3%) |
| Bevacizumab | 125 (44.0%) |
| OS (Months) |  |
| Median | 38 |
| IQR | 23.3-74.6 |

The DECIDER cohort includes tubo-ovarian tumors, solid omental and peritoneal metastases, and malignant ascites, sampled at diagnosis, after NACT, and at relapse. All 600 fresh-frozen samples underwent whole-genome sequencing (WGS) and RNA sequencing (RNA-seq), with DNA methylation additionally profiled by whole-genome bisulfite sequencing (WGBS) in 146 samples (Figure 1A). To link molecular findings with tissue architecture, we also analyzed 234 hematoxylin- and eosin-stained (H&E) whole-slide images from 57 patients.

**Figure 1.**
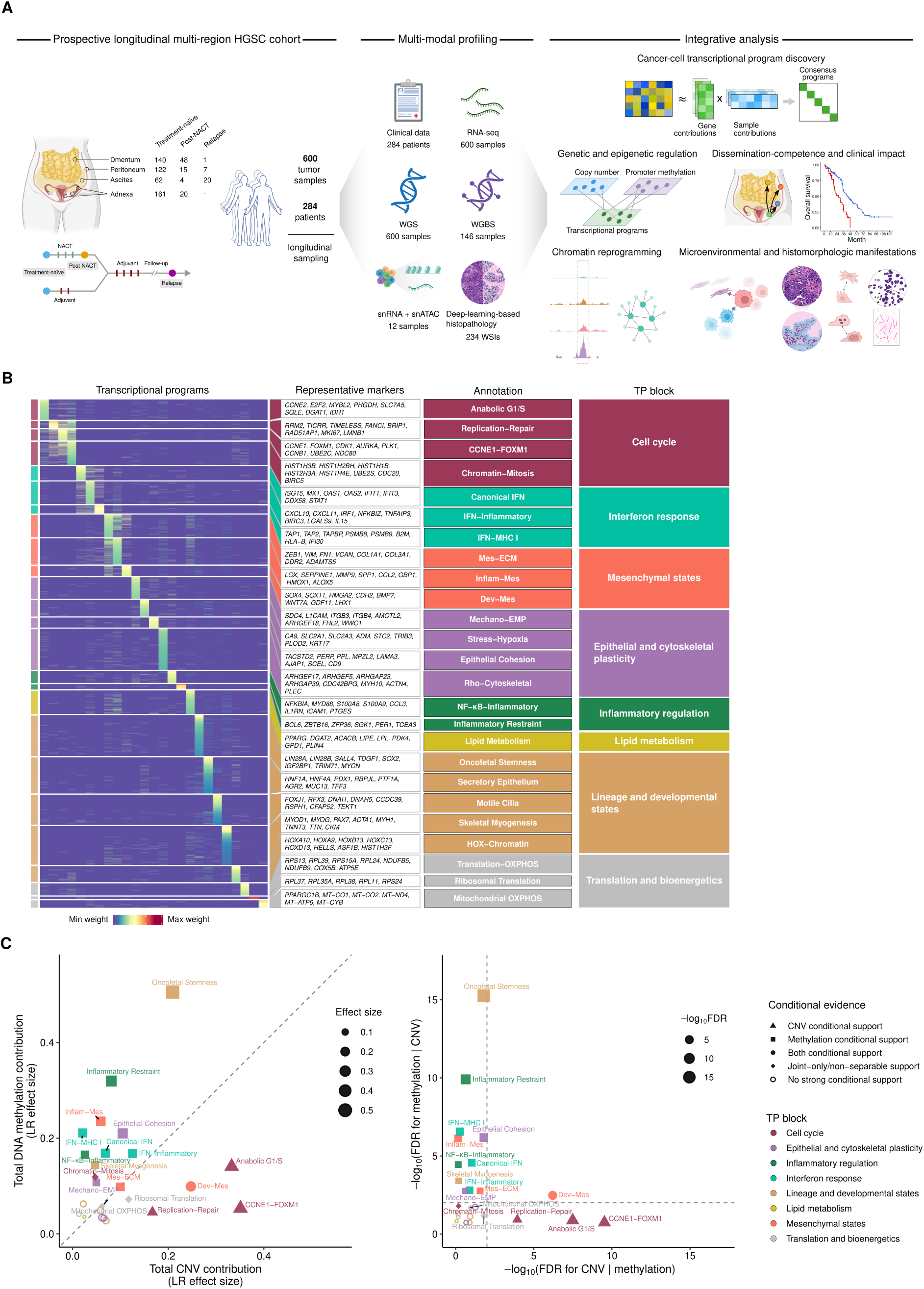
Study overview and characterization of transcriptional programs in HGSC. **(A)** Overview of the prospective longitudinal multi-region HGSC cohort, multi-modal molecular and histopathological profiling, and integrative analytical framework. “Adnexa” denotes the tubo-ovarian sampling site. **(B)** Heatmap shows normalized gene weights across the 25 identified transcriptional programs. Representative markers and biological annotations are shown alongside each program. **(C)** Genomic and epigenomic contributions to program activity in 146 treatment-naïve tumors from 85 patients with matched gene-expression, CNV and DNA-methylation profiles. Contributions and statistical support were estimated using likelihood-ratio comparisons of linear mixed-effects models accounting for tissue site and patient identity. Left, total contributions; point size denotes the joint CNV/methylation effect size, and the diagonal indicates equal contributions. Right, −log₁₀(FDR) for CNV and DNA-methylation associations after mutual adjustment; dashed lines indicate FDR = 0.01. Colors denote program blocks and shapes indicate conditional-evidence categories.

To resolve cancer-cell transcriptional states across tumors with varying purity, we used PRISM ^14^ to deconvolute bulk RNA-seq data into cancer-cell-specific expression profiles. We then applied consensus independent component analysis (ICA) to identify robust transcriptional programs (Methods). This approach revealed 25 recurrent programs spanning eight functional groups: cell cycle, epithelial and cytoskeletal plasticity, inflammatory regulation, interferon response, lineage and developmental states, lipid metabolism, mesenchymal states, and translation and bioenergetics (Figures 1B and S1A–S1C; Tables S1 and S2).

To investigate the genomic and epigenomic basis of transcriptional-program activity, we integrated program scores with matched copy number variation (CNV) and DNA methylation profiles. The strongest CNV associations were observed for three cell-cycle programs, i.e., Replication-Repair, CCNE1-FOXM1 and Anabolic G1/S (Figure 1C). Replication-Repair was associated with 15q gains encompassing *KIF23*, *TICRR* and *FANCI* (Figure S2A), CCNE1-FOXM1 with 19q12 amplification containing *CCNE1* (Figure S2B), and Anabolic G1/S with broad 8q24 gains (Figure S2C).

DNA methylation associations were strongest for Oncofetal Stemness and Inflammatory Restraint (Figure 1C). Their activities were inversely associated with methylation at developmental regulators including *SALL4* (Figure S2D) and antigen-presentation genes including *CIITA* and *CD74* (Figure S2E), respectively. Methylation was also associated with epithelial and mesenchymal plasticity programs. Mes-ECM activity was inversely associated with promoter methylation at extracellular-matrix and cytoskeletal genes, including *COL1A1* and *ACTA2* (Figure S2F), whereas Mechano-EMP showed inverse associations at genes involved in epithelial identity (*ELF3*), mechanotransduction (*FHL2*) and cellular stress responses (*CRYAB*) (Figure S2G). Dev-Mes was associated with both CNV and DNA methylation, with gene-level CNV associations at the 12q14.3 locus containing *HMGA2* and methylation associations at developmental and extracellular-matrix genes, including *LAMA1* and *WNT7A* (Figures S2H and S2I).

Together, these findings reveal distinct genomic and epigenomic associations across HGSC cancer-cell states. CNV is most strongly linked to proliferative programs, whereas DNA methylation is more closely associated with developmental, inflammatory and cell-state plasticity programs.

### Anatomical site defines transcriptional heterogeneity and chemotherapy response

Cancer cells disseminating within the peritoneal cavity encounter distinct tissue environments that may shape their transcriptional states. To investigate the contribution of anatomical context to cancer-cell heterogeneity in HGSC, we first examined how the 25 transcriptional programs varied across anatomical sites and between patients in treatment-naïve tumors. We used linear mixed-effects models to distinguish site-associated from inter-patient variation (Methods).

The relative contributions of anatomical site and patient identity differed across transcriptional programs (Figure 2A). Most lineage and developmental programs, together with translation- and bioenergetics-associated programs, showed greater variation between patients than across anatomical sites. In contrast, Mes-ECM, Mechano-EMP and NF-κB-Inflammatory exhibited the strongest site-associated variation, which exceeded inter-patient variation for Mechano-EMP and NF-κB-Inflammatory. This indicates that anatomical heterogeneity is particularly pronounced in cancer-cell programs of mesenchymal and epithelial plasticity and inflammatory signaling.

**Figure 2.**
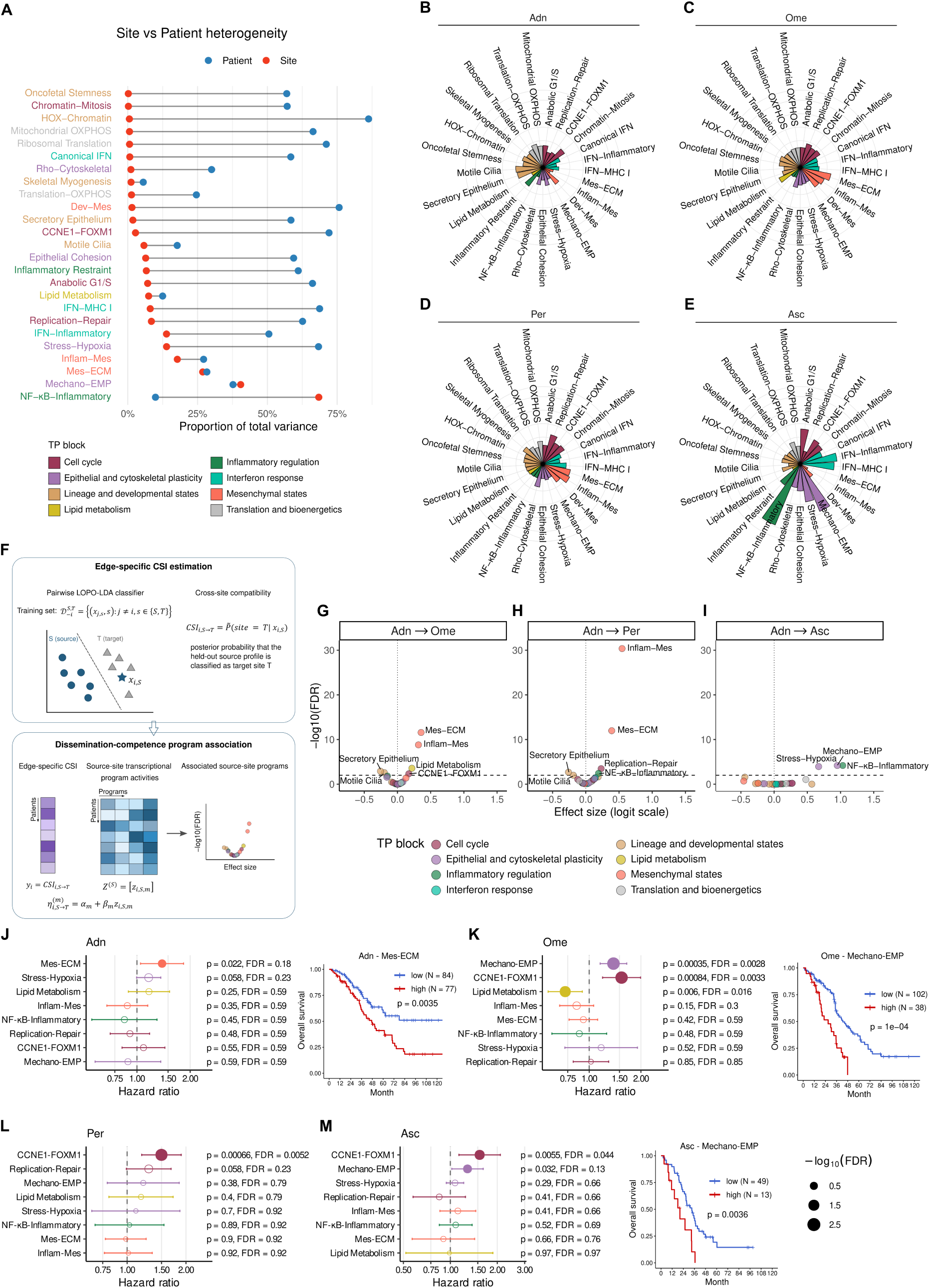
Anatomical site shapes transcriptional heterogeneity and reveals dissemination-associated programs with site-specific prognostic relevance. **(A)** Relative contributions of anatomical site and patient identity to variation in transcriptional-program activity across treatment-naïve tumors. For each program, red and blue points indicate the proportions of total variance attributable to anatomical site and patient identity, respectively, with connecting lines linking the two estimates. Program names are colored by program class. **(B–E)** Transcriptional-program activity across anatomical sites, shown separately for tubo-ovarian tumors (Adn; **B**), omental metastases (Ome; **C**), peritoneal metastases (Per; **D**), and malignant ascites (Asc; **E**). Radial bar lengths represent the site-specific estimated marginal mean activity of each program on a common scale, and bar colors denote program class. **(F)** Schematic overview of the cross-site inference (CSI) framework and dissemination-competence program analysis. For each specified source–target anatomical pair, CSI quantifies the extent to which the transcriptional profile of a source-site tumor resembles the profile characteristic of the target site. Source-site program activities are tested for association with CSI to identify programs linked to transcriptional compatibility with each target site. **(G–I)** Associations between transcriptional program activity in tubo-ovarian tumors and CSI for omental metastases **(G)**, peritoneal metastases (**H**), and ascites **(I)**. Each point represents one transcriptional program and is colored by program class. **(J–M)** Associations between pretreatment activity of dissemination-associated transcriptional programs and overall survival, assessed separately for tubo-ovarian tumors **(J)**, omental metastases **(K)**, peritoneal metastases **(L)** and malignant ascites **(M)**. Models were adjusted for treatment strategy. In the forest plots, point colors denote program class and point sizes represent −log10(FDR). Filled points indicate associations with nominal P≤0.05 and FDR ≤0.20; nominal P values and FDRs are shown alongside each program. Kaplan–Meier plots illustrate the site-specific associations for Mes-ECM in tubo-ovarian tumors **(J)**, Mechano-EMP in omental tumors **(K)**, and Mechano-EMP in ascitic tumors **(M)**.

We then examined how transcriptional-program activity differed across anatomical sites. Tubo-ovarian tumors showed higher activity of Secretory Epithelium and Motile Cilia programs, consistent with retention of fallopian tube-like epithelial features (Figures 2B and S3A). Solid omental and peritoneal metastases exhibited increased activity of mesenchymal programs, including Dev-Mes, Inflam-Mes and Mes-ECM, alongside Replication-Repair (Figures 2C, 2D, and S3A). In contrast, ascites showed higher activity of Mechano-EMP and Stress-Hypoxia, together with proliferative (CCNE1-FOXM1 and Anabolic G1/S) and inflammatory programs (IFN-Inflammatory, IFN-MHC I and NF-κB-Inflammatory; Figures 2E and S3A). These patterns reveal distinct anatomical distributions of cancer-cell states in treatment-naïve tumors, with mesenchymal programs predominating in solid metastases and epithelial-plasticity and stress–inflammatory programs in ascites.

We next asked whether CNV or DNA methylation could account for the anatomical variation in these programs, focusing on Mes-ECM, Mechano-EMP and NF-κB-Inflammatory, the three programs with the largest site-associated variance. In treatment-naïve tumors with matched molecular profiles, adjustment for DNA methylation reduced the site-associated variance of Mes-ECM by approximately 40%, with joint CNV–methylation adjustment increasing the reduction to approximately 45%. CNV adjustment alone resulted in no attenuation. In contrast, more than 90% of the site-associated variance in Mechano-EMP and NF-κB-Inflammatory remained after adjustment for both layers (Figure S3B). Thus, DNA methylation accounted for part of the anatomical variation in Mes-ECM, whereas the site dependence of Mechano-EMP and NF-κB-Inflammatory remained largely unexplained by measured CNV and DNA methylation, suggesting additional regulatory influences, potentially arising from the local microenvironment.

Chemotherapy-associated changes in cancer-cell transcriptional programs differed across anatomical sites. Relative to treatment-naïve samples from the corresponding sites, solid tumors collected after NACT showed reduced activity of proliferative programs, including Replication-Repair, Chromatin-Mitosis and CCNE1-FOXM1, alongside increased Epithelial Cohesion and Inflammatory Restraint (Figures S4A–S4C). In contrast, ascites showed markedly attenuated transcriptional changes, with significantly smaller chemotherapy-associated differences across this response pattern than those observed in solid tumors (Figures S4D and S4E). Comparisons between relapsed and treatment-naïve tumors revealed no significant within-site changes in program activity (Figure S4F). Together, these findings indicate that anatomical site is associated with both treatment-naïve transcriptional states and the magnitude of chemotherapy-associated remodeling, with pronounced changes in solid tumors but markedly attenuated changes in persistent ascites.

### Distinct transcriptional programs define competence for dissemination to solid metastatic sites and ascites

The distinct transcriptional states observed across solid metastases and ascites raised the possibility that corresponding features are already present in treatment-naïve tubo-ovarian tumors. To investigate this, we developed a Cross-Site Inference (CSI) framework that measures how closely the transcriptional profile of a source-site tumor resembles those of a candidate target site in other patients (Figure 2F; Methods). By evaluating individual source–target site pairs, CSI provided a patient-level measure of transcriptional compatibility with different metastatic compartments, allowing us to identify programs associated with site-biased dissemination.

CSI revealed distinct transcriptional programs associated with compatibility toward solid metastatic sites and ascites. Among the eight programs positively associated with at least one metastatic compartment, Mes-ECM and Inflam-Mes showed the strongest associations with both omental and peritoneal compatibility (Figures 2G and 2H). Lipid Metabolism and CCNE1-FOXM1 were additionally associated with omental compatibility, whereas Replication-Repair and NF-κB-Inflammatory were associated with peritoneal compatibility.

Ascites compatibility was associated with Mechano-EMP, NF-κB-Inflammatory and Stress-Hypoxia, reflecting a distinct combination of epithelial plasticity, inflammatory signaling and stress adaptation (Figure 2I). Together, these findings identify two broad patterns of metastatic-site compatibility already represented in tubo-ovarian tumors: a mesenchymal–stromal profile associated with solid metastatic tissues and a mechanoadaptive epithelial-plasticity and stress–inflammatory profile associated with ascites.

### Dissemination-associated programs stratify survival by anatomical site

To assess the clinical relevance of dissemination-associated transcriptional programs, we examined associations between their pretreatment activity and overall survival across anatomical sites using site-specific Cox proportional hazards models (Figures 2J–2M). Higher CCNE1-FOXM1 activity was associated with shorter survival across all three metastatic sites, but not in tubo-ovarian tumors. In contrast, higher Lipid Metabolism activity was associated with longer survival specifically in omental tumors.

Mes-ECM and Mechano-EMP showed particularly distinct site-specific associations. Higher Mes-ECM activity was associated with shorter survival in tubo-ovarian tumors, but not at metastatic sites. Conversely, higher Mechano-EMP activity was associated with shorter survival in omental tumors, with a similar trend in ascites, but showed no significant association in tubo-ovarian or peritoneal tumors (Figures 2J–2M).

The survival associations of Mes-ECM in tubo-ovarian tumors and Mechano-EMP in omental tumors remained significant after adjustment for age, treatment strategy, FIGO stage, residual disease after cytoreductive surgery and homologous recombination deficiency status (Figure S5). Together, the CSI and survival analyses link Mes-ECM and Mechano-EMP to distinct patterns of metastatic-site compatibility and reveal that their prognostic associations vary across anatomical compartments.

### A shared TEAD–AP-1 mechanotransductive core underlies distinct Mes-ECM and Mechano-EMP regulatory modes

The distinct metastatic-site associations of Mes-ECM and Mechano-EMP raise the question of whether these programs share underlying regulatory mechanisms. To investigate this, we performed joint single-nucleus RNA and ATAC profiling of 12 samples, yielding 79,279 high-quality nuclei, including 58,756 malignant epithelial, 6,826 stromal and 13,697 immune cells (Figures S6A–S6C). Based on Mes-ECM and Mechano-EMP activity, malignant epithelial cells were classified into four states: dual-low, Mes-ECM-high, Mechano-EMP-high and dual-high (Figure 3A).

**Figure 3.**
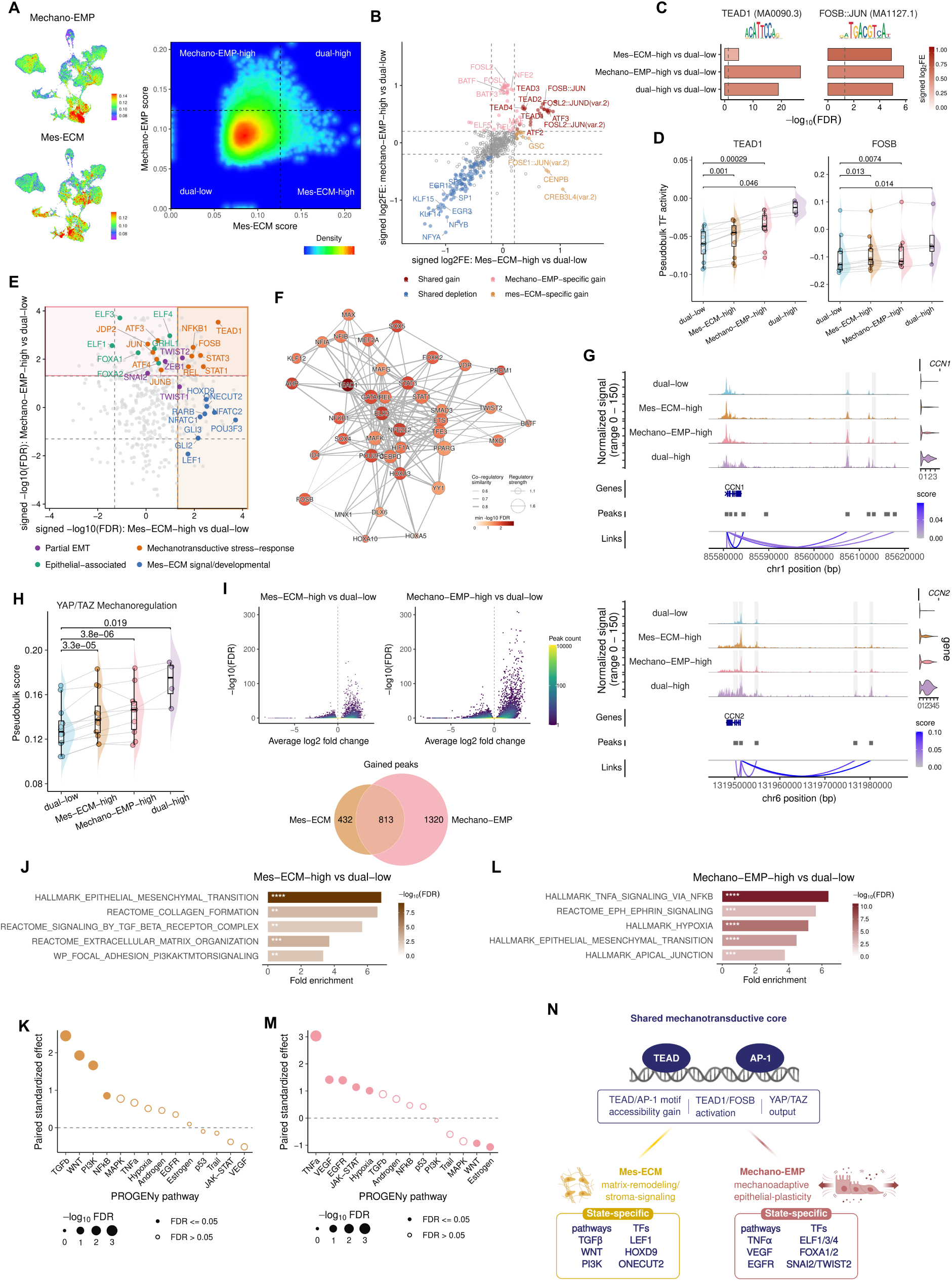
Shared TEAD–AP-1 mechanotransduction and distinct regulatory programs in Mes-ECM and Mechano-EMP cells. **(A)** UMAPs of cancer cells colored by Mechano-EMP or Mes-ECM score, and a two-dimensional density plot of Mes-ECM and Mechano-EMP scores. Dashed lines indicate the thresholds used to define dual-low, Mes-ECM-high, Mechano-EMP-high, and dual-high cells. **(B)** Comparative motif enrichment among differentially accessible peaks in Mes-ECM-high versus dual-low and Mechano-EMP-high versus dual-low cells. Each point represents a motif. Axes show signed log2 fold enrichment, with positive and negative values indicating enrichment among gained and depleted peaks, respectively. Dashed lines indicate ∣log2FE∣=0.2. Motifs meeting FDR < 0.05 and ∣log2FE∣>0.2 in the relevant comparison or comparisons were classified as shared gain, shared depletion, or state-specific gain motifs. **(C)** Bar plots showing motif enrichment for exemplary TFs: TEAD1 and FOSB::JUN motifs in Mes-ECM-high, Mechano-EMP-high, and dual-high cells relative to dual-low cells. Bar length indicates −log10(FDR), and fill indicates signed log2 fold enrichment. **(D)** Sample–state pseudobulk activities of TEAD1 and FOSB across the four malignant-cell states. Points denote sample–state pseudobulks, and lines connect matched states within the same sample. Each non-dual-low state was compared with the matched dual-low state using a two-sided paired t-test. P values were adjusted across all TF–contrast tests using the Benjamini–Hochberg method. **(E)** Comparative TF-activity landscape for Mes-ECM-high versus dual-low and Mechano-EMP-high versus dual-low cells. Axes show signed −log10(FDR), and dashed lines indicate FDR = 0.05. Selected epithelial-associated, partial-EMT, mechanotransductive stress-response, and Mes-ECM signaling/developmental regulators are highlighted. Statistical testing and multiple-testing correction were performed as in **(D)**. **(F)** TF–TF co-regulatory similarity network of TFs with significantly increased activity in both Mes-ECM-high and Mechano-EMP-high cells relative to dual-low cells (FDR < 0.05 in both comparisons). Node fill indicates the minimum −log10(FDR) across the two comparisons, node size indicates the log-transformed sum of retained positive TF–target regulatory weights, and edge width indicates co-regulatory similarity. **(G)** Aggregated chromatin-accessibility profiles, peak-to-gene links, and gene-expression distributions for the canonical YAP/TEAD targets *CCN1* and *CCN2* across the four malignant-cell states. **(H)** Sample–state pseudobulk scores for a YAP/TAZ mechanoregulation gene set across the four cancer-cell states. Points denote pseudobulks, and lines connect matched states within the same sample. Each non-dual-low state was compared with the matched dual-low state using a two-sided paired t-test. Exact two-sided P values are shown. **(I)** Differential chromatin accessibility in Mes-ECM-high versus dual-low and Mechano-EMP-high versus dual-low cells. Hexagon-binned volcano plots show average log2 fold change and −log10(FDR) across accessible regions. The Venn diagram shows the overlap between gained peaks in the two comparisons. **(J)** Representative gene sets enriched among upregulated genes positively linked to peaks with increased accessibility in Mes-ECM-high versus dual-low cells. Bar length indicates fold enrichment, and fill indicates −log10(FDR). Asterisks indicate FDR thresholds: ∗FDR < 0.05, ∗∗FDR < 0.01, ∗∗∗FDR < 0.001, and ∗∗∗∗FDR < 0.0001. **(K)** Differential PROGENy pathway activity in Mes-ECM-high versus dual-low cells. Each point represents a pathway; the vertical axis shows the paired standardized effect size calculated from the matched sample–state pseudobulk activity differences. Point size indicates −log10(FDR); filled and open points denote FDR ≤ 0.05 and > 0.05, respectively. **(L)** Representative gene sets enriched among upregulated genes positively linked to peaks with increased accessibility in Mechano-EMP-high versus dual-low cells. Bar length indicates fold enrichment, and fill indicates −log10(FDR). Asterisks denote FDR thresholds as in **(J)**. **(M)** Differential PROGENy pathway activity in Mechano-EMP-high versus dual-low cells. Plotting conventions and statistical analysis are as in **(K)**. **(N)** Schematic summarizing regulatory programs in Mes-ECM and Mechano-EMP states.

Despite their distinct anatomical associations, Mes-ECM-high and Mechano-EMP-high cells shared a prominent regulatory signature characterized by increased chromatin accessibility at TEAD and AP-1 motifs (Figures 3B and 3C). Consistently, TEAD1 and FOSB regulon activities were elevated in both states and in dual-high cells (Figure 3D). Co-regulatory network analysis further linked TEAD1 and AP-1-related factors with inflammatory, hypoxia-responsive and other signal-responsive regulators (Figures 3E and 3F). This shared TEAD–AP-1 signature was accompanied by increased chromatin accessibility and expression of the canonical YAP/TEAD targets *CCN1* and *CCN2*, together with elevated YAP/TAZ mechanoregulation activity (Figures 3G and 3H). Together, these findings identify a shared TEAD–AP-1-associated mechanotransductive regulatory program across the two plasticity states.

Mes-ECM-high cells showed a regulatory profile dominated by the shared TEAD–AP-1 signature, with most gains in chromatin accessibility also observed in Mechano-EMP-high cells and relatively few state-specific enriched motifs (Figures 3B and 3I). Transcription-factor activity revealed preferential activation of developmental and signal-responsive regulators, including LEF1 and NFAT-family factors (Figure 3E). Upregulated genes associated with increased chromatin accessibility were enriched for extracellular-matrix organization, collagen formation, TGFβ-receptor signaling and focal-adhesion–PI3K–AKT–mTOR signaling (Figure 3J), supported by pathway-activity analysis showing elevated TGFβ, WNT and PI3K activity (Figure 3K). Together, these findings link the shared mechanotransductive program in Mes-ECM-high cells to a distinct regulatory profile associated with extracellular-matrix remodeling and cell–matrix adhesion.

By contrast, Mechano-EMP-high cells exhibited more extensive chromatin remodeling, with state-specific enrichment of AP-1/BATF and REL-family motifs beyond the shared TEAD–AP-1 signature (Figures 3B and 3I). Transcription-factor activity analysis revealed concurrent activation of epithelial-associated regulators (FOXA1/2 and ELF1/3/4), partial EMT regulators (SNAI2 and TWIST2) and stress-responsive factors (Figure 3E). Consistent with this regulatory profile, upregulated genes linked to gained chromatin-accessibility peaks were enriched for TNFα–NF-κB signaling, hypoxia, EMT, apical-junction remodeling and ephrin signaling (Figure 3L). Pathway analysis further revealed increased EGFR and JAK–STAT activity, alongside TNFα signaling and hypoxia (Figure 3M). Mechano-EMP thus represents a hybrid epithelial–mesenchymal state that integrates inflammatory and stress-adaptive regulation with the shared mechanotransductive core (Figure 3N).

Dual-high cells combined the matrix-remodeling programs of Mes-ECM with the inflammatory and stress-adaptive programs of Mechano-EMP (Figure S7). This composite state integrates mesenchymal remodeling and mechanoadaptive epithelial plasticity within a shared TEAD–AP-1-centered regulatory landscape.

### Mes-ECM and Mechano-EMP engage distinct tumor–microenvironment interactions

The distinct regulatory features of Mes-ECM and Mechano-EMP suggested that the two cancer-cell states engage their surrounding microenvironments through different signaling interactions. We therefore used ligand–receptor inference to examine their communication with 14 TME cell types spanning fibroblastic, vascular, adipocytic, myeloid, lymphoid and dendritic populations (Figures 4A and 4B).

**Figure 4.**
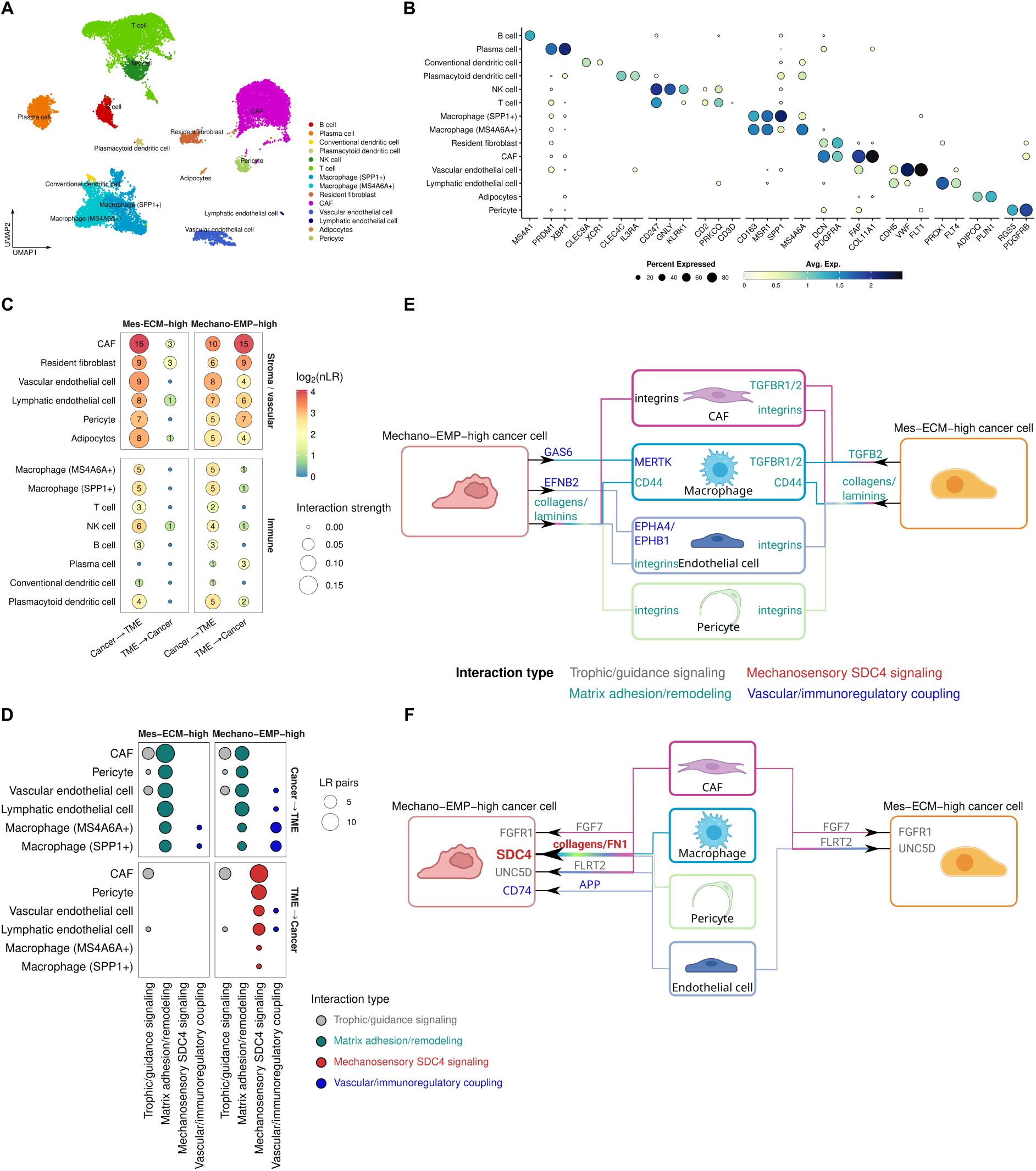
Distinct tumor–microenvironment interactions characterize Mes-ECM and Mechano-EMP cancer states. **(A)** UMAP of tumor-microenvironment (TME) cells colored by annotated cell type. **(B)** Expression of representative marker genes across the annotated TME populations. Dot size indicates the percentage of cells expressing each gene and color indicates average expression. **(C)** Number and aggregate strength of differentially detected ligand–receptor interactions between Mes-ECM-high or Mechano-EMP-high compared to dual-low cancer cells and TME populations, shown separately for cancer-to-TME and TME-to-cancer directions. Numbers within circles indicate the number of ligand–receptor pairs, circle color indicates log2-transformed interaction number, and circle size indicates summed interaction strength. **(D)** Categorization of differentially detected ligand–receptor interactions between Mes-ECM-high or Mechano-EMP-high compared to dual-low cancer cells and the indicated TME populations, shown separately for cancer-to-TME and TME-to-cancer directions. Interactions are grouped as trophic/guidance signaling, matrix adhesion/remodeling, mechanosensory SDC4 signaling or vascular/immunoregulatory coupling; circle size indicates the number of ligand–receptor pairs. **(E–F)** Schematics of major cancer-to-TME **(E)** and TME-to-cancer **(F)** interactions involving cancer-associated fibroblasts (CAFs), macrophages, endothelial cells and pericytes. Gene-label colors indicate interaction categories.

Ligand–receptor analysis revealed distinct communication patterns for Mes-ECM-high and Mechano-EMP-high cancer cells relative to dual-low cells. Both states shared predicted inputs from cancer-associated fibroblasts (CAFs), including FGF7–FGFR1 and FLRT2–UNC5D signaling (Figure S8A). Beyond these shared inputs, the two states differed markedly in the direction and cellular origin of microenvironmental signaling: Mes-ECM-high cells showed a strong bias toward outgoing cancer-to-microenvironment interactions, whereas Mechano-EMP-high cells received prominent inputs from multiple stromal, vascular and macrophage populations (Figures 4C and 4D).

Mes-ECM-high cells showed predominantly outgoing matrix-adhesion and guidance signals toward CAFs, pericytes, endothelial cells and macrophages, together with TGFB2–TGFBR1/2 signaling toward CAFs and macrophages (Figures 4E and S8B–S8E). Reciprocal inputs from pericytes, endothelial cells and macrophages were limited (Figure 4C). Mechano-EMP-high cells also exhibited outgoing interactions with vascular and myeloid populations, including EFNB2–EPH signaling toward endothelial cells and GAS6–MERTK and MIF–CD74/CD44 signaling toward macrophages (Figures 4E, S8C, and S8E).

A defining feature of Mechano-EMP-high cells was the convergence of microenvironmental signals on SDC4, a mechanosensory co-receptor involved in integrin signaling and force transduction. CAFs, pericytes and endothelial cells contributed collagen-, fibronectin- and thrombospondin-mediated inputs to SDC4 (Figures 4D, 4F, S8A, S8F, and S8G). Additional FN1–SDC4 interactions originated from macrophages, particularly SPP1+ macrophages (Figure S8H). These SDC4-centered inputs were not detected for Mes-ECM-high cells. Mechano-EMP-high cells also received endothelial APP–CD74 signals (Figures 4F and S8F). This distinct communication pattern shows that Mes-ECM is associated primarily with outward signaling toward the surrounding stroma, whereas Mechano-EMP is characterized by a convergent stromal–vascular–myeloid input network that combines SDC4-centered matrix sensing with inflammatory signaling.

### Histomorphologic manifestations of dissemination-associated tumor states

To assess whether Mes-ECM and Mechano-EMP are reflected in tissue architecture, we analyzed 234 H&E-stained whole-slide images from 57 patients using deep-learning-based panoptic segmentation ^15^ and quantitative assessment of stromal, extracellular-matrix and cancer-cell features (Figure 5A). To distinguish the associations of each program, we compared Mes-ECM-high and Mechano-EMP-high tumors, with the other program held low, against tumors with low activity of both programs using linear mixed-effects models.

**Figure 5.**
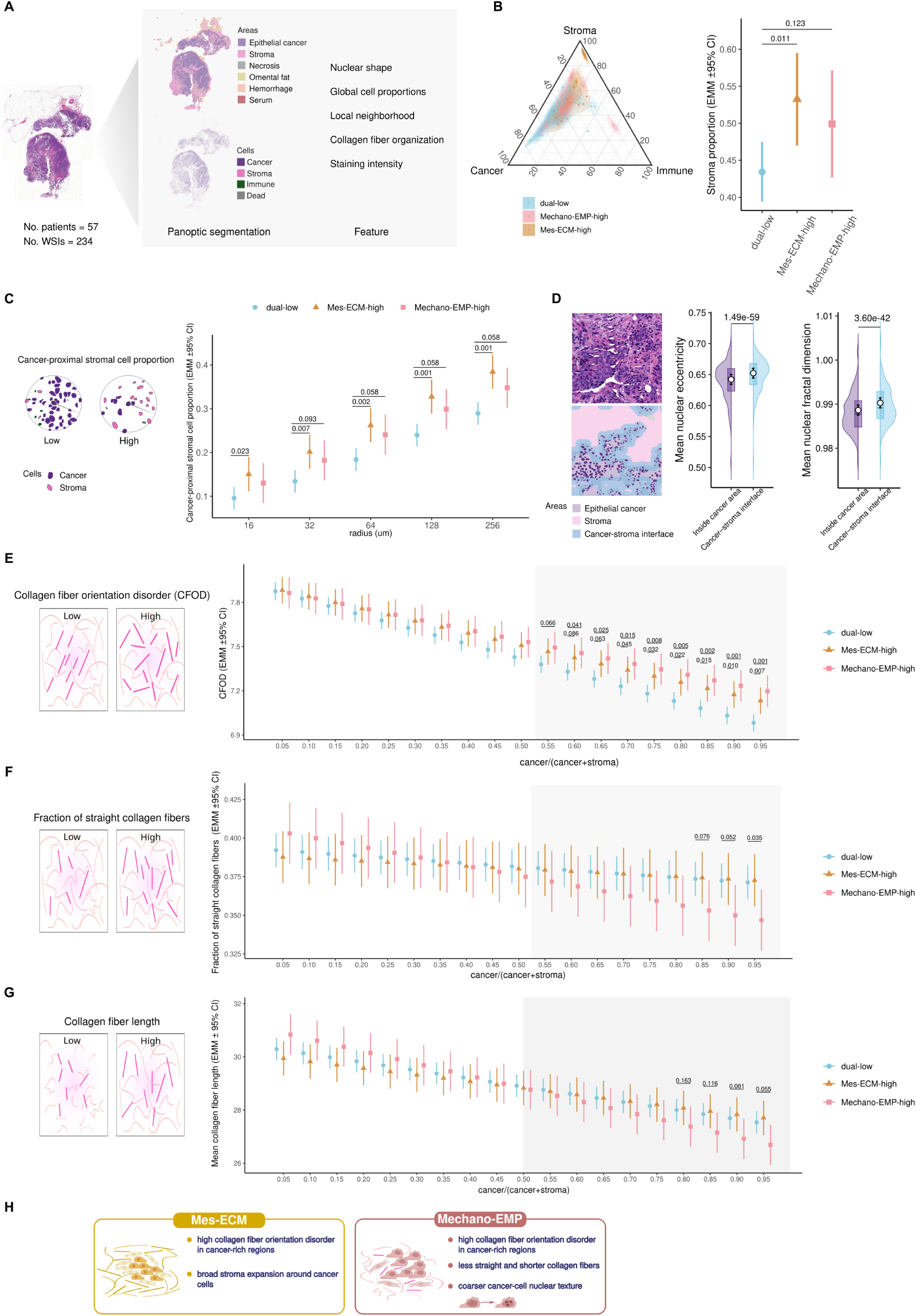
Histomorphologic manifestations of dissemination-associated tumor states. **(A)** Overview of panoptic segmentation and quantitative histomorphologic analysis of H&E whole-slide images. **(B)** Global cellular composition. Left, ternary plot of relative cancer, stromal and immune cell proportions across transcriptional states. Right, estimated marginal mean (EMM) stromal-cell proportions with 95% confidence intervals (CIs). **(C)** Cancer-proximal stromal-cell proportions within cancer-centered neighborhoods of increasing radius. Points and error bars show EMMs and 95% CIs. Benjamini–Hochberg-adjusted P values < 0.1 are shown. **(D)** Cancer-cell nuclear morphology at the cancer–stroma interface. Left, representative segmentation distinguishing cancer cells within a 16-µm interface region from those within the tumor interior. Middle and right, distributions of mean nuclear eccentricity and fractal dimension at the interface and within the tumor interior, overlaid with EMMs and 95% CIs. P values indicate two-sided paired within-slide contrasts between regions. **(E–G)** Collagen-fiber orientation disorder (CFOD; **E**), fraction of straight collagen fibers **(F)** and mean collagen-fiber length **(G)** across local cancer–stroma compositions. Schematics illustrate each feature; plots show EMMs and 95% CIs across the local cancer fraction, defined as cancer/(cancer + stroma). Shading marks cancer-rich compositions. Higher CFOD indicates greater orientation disorder. P values < 0.1 are shown in **(E)** and **(F)**, and P values < 0.2 in **(G)**. **(H)** Summary of state-associated histomorphologic features. Estimates were obtained from linear mixed-effects models. Mes-ECM-high and Mechano-EMP-high denote high activity of the indicated program with the other held low; dual-low denotes low activity of both programs.

Mes-ECM was associated with global and cancer-proximal stromal expansion. Mes-ECM-high tumors showed an increased global stromal-cell proportion (P = 0.011), whereas the corresponding increase in Mechano-EMP-high tumors was not significant (P = 0.12; Figure 5B). Spatial analysis further revealed increased stromal-cell proportions in the immediate vicinity of cancer cells in Mes-ECM-high tumors (neighborhood radii of 16–32 µm; P = 0.007–0.023), extending to broader neighborhoods (64–256 µm; P = 0.001–0.002; Figure 5C). Mechano-EMP-high tumors showed only modest trends in the same direction. These findings identify global and cancer-proximal stromal accumulation as prominent tissue features of Mes-ECM-high tumors. Cancer cells at the tumor–stroma interface also exhibited greater nuclear eccentricity and fractal dimension than those in tumor interiors, revealing morphological differences associated with local stromal proximity (Figure 5D).

Both Mes-ECM-high and Mechano-EMP-high tumors exhibited increased collagen orientation disorder within cancer-rich regions, whereas no corresponding differences were observed in stroma-rich regions (Figures 5E and S9). This composition-dependent pattern suggests that altered collagen organization is associated with the local cancer-cell microenvironment rather than simply reflecting differences in overall stromal abundance.

Additional differences emerged in collagen fiber morphology. Mechano-EMP-high tumors showed trends toward shorter and less straight collagen fibers in cancer-rich stromal regions, while comparable changes were not observed in Mes-ECM-high tumors (Figures 5F and 5G). These patterns suggest that Mechano-EMP is associated with additional localized alterations in matrix organization.

Mechano-EMP-high tumors also displayed altered cancer-cell nuclear texture, characterized by fewer but larger hematoxylin-dense chromatin patches compared with dual-low tumors (Figure S10). This coarser nuclear texture provides a further morphological correlate of the Mechano-EMP transcriptional state.

The histomorphological analyses reveal distinct but partly overlapping tissue features associated with the two plasticity programs. Mes-ECM is characterized by extensive cancer-proximal stromal accumulation, whereas Mechano-EMP shows additional associations with localized collagen fiber alterations and coarser cancer-cell nuclear texture. Both programs share increased collagen orientation disorder in cancer-rich regions (Figure 5H). These tissue-level patterns extend the molecular and microenvironmental organization of Mes-ECM and Mechano-EMP states to spatially resolved histomorphological phenotypes.

## DISCUSSION

Intraperitoneal dissemination exposes cancer cells to fundamentally different environments, requiring persistence in peritoneal fluid and colonization of solid tissues. Our study defines a site-resolved transcriptional landscape of HGSC and identifies distinct configurations of cancer-cell plasticity associated with these metastatic compartments. Previous studies have established the importance of mesenchymal programs for mesothelial clearance, inflammatory signaling in malignant ascites, and anatomical heterogeneity in HGSC ^12,13,16,17^. We extend this understanding by linking recurrent cancer-cell programs already represented in tubo-ovarian tumors to distinct solid-metastatic and ascitic transcriptional phenotypes. These observations support a model in which intraperitoneal dissemination is associated with anatomically specialized cancer-cell states rather than a single, universal metastatic program.

This perspective complements genomic studies that have established the evolutionary complexity and non-random dissemination of HGSC ^7–9^. Although genomic alterations contribute to transcriptional heterogeneity, our analyses showed that the anatomical distributions of several plasticity programs were largely unexplained by measured CNV and DNA methylation. The distinct molecular associations of Mes-ECM and Mechano-EMP further highlight that metastatic cell states can differ in their relationships with intrinsic molecular variation and anatomical context. Recent studies in colorectal and pancreatic cancer similarly implicate transcriptional plasticity and microenvironmental influences in metastatic adaptation ^18,19^. Our findings extend this perspective to transcoelomic dissemination, where solid metastases and malignant ascites coexist within the same disease.

A central insight is that anatomically distinct plasticity states share a common mechanotransductive regulatory architecture. TEAD–AP-1 cooperation and YAP/TAZ signaling are established regulators of cancer-cell behavior ^20^, but their association with different metastatic phenotypes in HGSC suggests a broader role for this regulatory axis. Mes-ECM and Mechano-EMP couple the shared mechanotransductive program to different regulatory and microenvironmental contexts, favoring ECM–stromal remodeling and stress–inflammatory responsiveness, respectively. The convergence of stromal, vascular and myeloid inputs on SDC4 in Mechano-EMP is particularly intriguing given the established role of SDC4 in force-dependent integrin–RhoA–YAP signaling ^21^. These interactions nominate a potential connection between microenvironmental signals and cancer-cell mechanoregulation. The coexistence of both programs in dual-high cells further supports a modular organization of plasticity, in which distinct regulatory features can operate within a shared cellular framework rather than defining mutually exclusive states.

The histomorphological associations provide a tissue-level dimension. Mes-ECM was associated with cancer-proximal stromal accumulation, whereas both programs showed altered collagen organization within cancer-rich regions. These patterns suggest that transcriptional plasticity is reflected in the spatial architecture of the tumor microenvironment, providing candidate accessible morphological markers of these two cancer-cell plasticity states.

The anatomical context of cancer-cell states has important clinical implications. The site-specific prognostic associations and differences in chemotherapy-associated transcriptional remodeling indicate that the biological and clinical significance of a cancer-cell program cannot be interpreted independently of where it is expressed. The attenuated remodeling observed in persistent ascites extends previous evidence of compartment-specific treatment responses in HGSC ^13^. These observations emphasize the importance of multisite sampling for biomarker development and therapeutic stratification. The convergence of Mes-ECM and Mechano-EMP on a shared mechanotransductive signature nominates this axis as a common therapeutic entry point across solid metastatic and ascitic disease, whereas their divergent ECM–stromal and stress–inflammatory branches provide a rationale for state-specific combination strategies. More broadly, our study positions cancer-cell state, anatomical context and microenvironmental interactions as interconnected dimensions of metastatic heterogeneity that should be considered together when investigating and targeting disseminated cancer.

## Limitations of the study

Several limitations should be considered. CSI identifies transcriptional compatibility between anatomical sites but cannot establish metastatic seeding capacity, reconstruct dissemination events or determine how Mes-ECM and Mechano-EMP states evolve during metastatic progression. The attenuated chemotherapy-associated transcriptional changes observed in post-NACT ascites may reflect either limited remodeling of persisting cancer cells or selection of treatment-tolerant populations. Because post-NACT ascites was available only from patients with residual fluid, these possibilities cannot be distinguished in the present cohort.

## METHODS

### Patient cohort

This study included 284 patients with HGSC enrolled in DECIDER, a prospective, longitudinal, multiregion observational trial (ClinicalTrials.gov identifier: NCT04846933). DECIDER enrolled patients older than 18 years who were treated for suspected ovarian cancer at Turku University Hospital, Finland, and were able and willing to provide written informed consent. No exclusion criteria were applied at enrollment; HGSC histology was required for the present study. All participants provided written informed consent. The study and use of clinical material were approved by the Ethics Committee of the Hospital District of Southwest Finland (decision number: EMTK: 145/1801/2015).

### RNA-seq data processing

Fresh tumor samples were collected during laparoscopy or debulking surgery, or from drained ascites. Samples meeting RNA quantity and quality criteria underwent library preparation and sequencing at BGI (BGI Europe A/S, Denmark) or Novogene (Novogene Europe, UK). RNA sequencing was performed on the BGISEQ-500, MGISEQ-2000, HiSeq 2000, HiSeq 4000, HiSeq X Ten or NovaSeq 6000 platform, generating 100- or 150-bp paired-end reads. Bulk RNA-seq data were processed using the SePIA pipeline ^22^ within the Anduril2 framework ^23^, as previously described ^14^. Read quality was assessed using FastQC v0.11.4 ^24^, and sequencing adapters and low-quality bases were trimmed using Trimmomatic v0.33 ^25^. Trimmed reads were aligned to the GRCh38.d1.vd1 reference genome with GENCODE v25 annotation using STAR v2.5.2b ^26^, allowing up to 10 mismatches per read and retaining all reported alignments. Gene-level effective counts were quantified using eXpress v1.5.1 ^27^. Cell-type proportions and cell-type-specific expression profiles were inferred from bulk expression profiles using PRISM ^14^.

### Whole genome sequencing data processing

Samples meeting DNA quantity and quality criteria underwent library preparation and sequencing at BGI or Novogene. Whole-genome sequencing (WGS) was performed on the BGISEQ-500, MGISEQ-2000, HiSeq X Ten or NovaSeq 6000 platform, generating 100- or 150-bp paired-end reads. WGS data were processed as previously described ^9^. Read quality was assessed using FastQC v0.11.4 ^24^, and sequencing adapters and low-quality bases were trimmed using Trimmomatic v0.33 ^25^. Trimmed reads were aligned to GRCh38.d1.vd1 using BWA-MEM v0.7.12-4103942 ^28^ with the -M option. Duplicate reads were removed using Picard v2.6 (https://github.com/broadinstitute/picard), and base quality scores were recalibrated using BaseRecalibrator ^29^ in the Genome Analysis Toolkit (GATK) v3.7 ^30^. We used GATK 4.1.4.1 ^30^ to perform copy-number segmentation and estimated purity, ploidy, and purified logR values using a reimplemented ASCAT algorithm ^31^ as described in ^32^.

### Whole genome bisulfite sequencing data processing

Tumor samples underwent bisulfite conversion, library preparation and whole-genome bisulfite sequencing (WGBS) at BGI (BGI Europe A/S, Denmark). WGBS data were processed as described in ^33^. Reads were trimmed using Trimmomatic v0.32 ^25^ and assessed for quality using FastQC v0.11.4 ^24^. Trimmed reads were aligned to GRCh38.d1.vd1 using Bismark v0.22.3 ^34^ with the --bowtie2 option, followed by deduplication using deduplicate_bismark. Methylation β-values were obtained using bismark_methylation_extractor for 58,304,913 CpG sites, with sites on the forward and reverse strands counted separately. We applied deconvolution to profile the cancer methylome across different tissues, patient groups, and anticancer treatments ^35^, allowing us to isolate the cancer-derived methylation signals, which were subsequently used for downstream analyses.

### Independent component analysis and derivation of transcriptional programs

#### Consensus ICA decomposition

ICA was selected as the module-detection framework based on a comprehensive benchmark of 42 gene-expression module-detection methods, in which ICA-based decomposition methods most consistently recovered known regulatory modules across datasets ^36^. Gene-wise biological variance was estimated from the log-normalized expression matrix using modelGeneVar from scran (version 1.26.2) ^37^. Genes with a nominal variance-model *P* < 0.2 and positive estimated biological variance (*bio* > 0) were retained, yielding 4,830 highly variable genes. The resulting sample-by-gene matrix was centered and scaled gene-wise before dimensionality reduction.

An upper bound for ICA dimensionality was determined using permutation-based parallel analysis of principal-component (PC) eigenvalues. Truncated principal component analysis (PCA) was used to calculate the first 150 observed eigenvalues. Null eigenvalue distributions were generated from 200 permuted expression matrices, in each of which expression values were independently shuffled across samples within each gene, preserving gene-specific expression distributions while disrupting covariance among genes. At each component rank, the observed eigenvalue was compared with the corresponding 95th percentile of the permutation distribution. The number of observed eigenvalues exceeding these rank-matched thresholds defined *K_max_*, yielding K_max_ = 81.

Candidate ICA solutions with K = 2, …, 81 were evaluated after projecting the standardized expression matrix onto 150 principal components. For each K, FastICA (version 1.2.7) ^38,39^ was run 100 times with random initializations using the log-cosh contrast function and deflationary component extraction. ICA mixing vectors estimated in PCA space were mapped back to the 4,830-gene expression space. For each candidate K, all gene-loading vectors were pooled and clustered into K groups using *k*-medoids with the sign-invariant dissimilarity 1 − |cor|. Component stability was defined as the mean pairwise absolute correlation among loading vectors within each cluster. Candidate solutions were summarized by their mean component stability and the fraction of components with stability ≥ 0.75. Among solutions with K > 15, K = 24 had the highest mean stability and fraction of stable components and was selected for the final decomposition (Figure S1A).

For K = 24, FastICA was repeated 300 times. The resulting 7,200 gene-loading vectors were clustered into 24 groups using the same correlation-based k-medoids procedure. Within each cluster, the medoid was used as an orientation reference: loading vectors were sign-aligned to the medoid and then averaged. The resulting 24 averaged vectors formed the consensus gene-loading matrix.

### Derivation of candidate transcriptional programs

To identify genes with significant ICA loadings, each consensus component was assigned a canonical orientation by requiring the summed loadings of the top 10% of genes ranked by absolute loading to be positive. For each component, a robust reference scale was estimated preferentially using the unscaled Q_n_ statistic calculated with robustbase::Qn(…, constant = 1), with central-IQR, non-zero-IQR, and standard-deviation fallbacks when required. Gene-level loading statistics were obtained by dividing each loading by the component-specific reference scale, and two-sided normal-reference P-values were calculated. P-values were adjusted using the Benjamini–Hochberg procedure, and genes with adjusted q ≤ 10^−5^ were considered to have significant loadings on that component.

Within each component, genes with significant positive and negative loadings were used to define separate candidate transcriptional programs. Programs containing fewer than 15 genes were excluded. Positive-loading programs retained their positive ICA loadings, whereas negative-loading programs used the absolute magnitudes of their negative loadings, such that higher downstream scores represented stronger expression of the genes defining each program. Each sparse candidate-program weight vector was L2-normalized to unit Euclidean norm, producing a weighted gene-by-program matrix that preserved relative ICA loading magnitudes within each program.

### Filtering of batch-associated components and programs

Sequencing-batch effects were assessed at the level of sample-specific ICA component scores. For each component, we fitted a linear mixed-effects model,

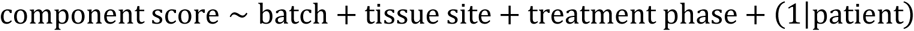

the overall batch effect was assessed using a likelihood-ratio test comparing maximum-likelihood fits of the full model and the corresponding model without batch. Batch effect size was quantified as the non-negative difference in Nakagawa marginal R2 between the full and reduced models. Components with *R^2^_batch_* and a batch-effect P < 0.01 were classified as substantially batch-associated, and all candidate transcriptional programs derived from these components were removed from the weighted program matrix.

#### Functional annotation of transcriptional programs

To functionally annotate the remaining candidate transcriptional programs, we performed gene-set over-representation analysis using the MSigDB Hallmark, Reactome, and WikiPathways collections ^40–43^. Gene sets were restricted to those containing 20–1,000 genes after intersection with the background gene universe. For each candidate program, enrichment was assessed using a one-sided hypergeometric test, with P-values adjusted across gene sets using the Benjamini–Hochberg procedure. Gene sets were considered significantly enriched at adjusted P < 0.01 with at least five overlapping program genes. Candidate programs lacking any gene set meeting these criteria were excluded from further analyses, resulting in 25 retained transcriptional programs. Significantly enriched gene sets were used to guide functional annotation and naming of the retained programs.

#### Estimation of sample-level transcriptional-program activity

Sample-level transcriptional-program scores were calculated from the final weight matrix after removal of programs derived from batch-associated components and programs lacking significant gene-set enrichment. Gene-expression values were centered and scaled across samples, and the activities of all retained programs were estimated jointly by ordinary least squares using a pivoted QR decomposition of the filtered weight matrix. Thus, for each sample, the standardized expression profile was modeled as a linear combination of the retained weighted transcriptional programs, with each program score estimated conditional on the other retained programs.

### Attribution of transcriptional-program activity to CNV and DNA methylation

Program-level CNV and DNA methylation associations were evaluated in 146 treatment-naïve tumors from 85 patients with matched RNA-sequencing, purified-logR CNV and DNA methylation M-value data. For each transcriptional program and molecular layer, analyses were restricted to features corresponding to program-member genes, which were standardized before principal-component analysis. For each molecular layer, we retained the minimum number of PCs explaining at least 30% of the cumulative variance to limit model complexity, with a maximum of ten PCs; this yielded three to six CNV PCs and two to four methylation PCs across programs.

For each program, four linear mixed-effects models were fitted by maximum likelihood with tissue site as a fixed effect and patient as a random intercept: a baseline model containing no molecular covariates (*M*_0_), CNV-only (*M_C_*) and methylation-only (*M_M_*) models containing the respective retained PCs, and a joint CNV–methylation model (M_CM_). Total CNV and methylation associations were quantified by comparing *M*_0_ with *M_C_* and M_M_, respectively, and the joint molecular association by comparing *M*_0_ with *M_CM_*. Modality-specific associations after mutual adjustment were quantified by comparing *M_M_* with *M_CM_* for CNV conditional on methylation and *M_C_* with *M_CM_* for methylation conditional on CNV. Likelihood-ratio statistics were transformed as E = 1 − exp(−LR/n), where *n* is the number of observations, to provide a likelihood-based measure of incremental model fit. Models were fitted using lmer in the lme4 R package ^44^ with the bobyqa optimizer, and likelihood-ratio comparisons were retained only when both nested models converged and used the same observations. *P* values from the likelihood-ratio tests were Benjamini–Hochberg adjusted across transcriptional programs. Using an FDR threshold of 0.01, programs were classified as showing CNV conditional, methylation conditional, both conditional, joint-only, or no strong support according to the significance pattern of the conditional and joint tests.

Gene-level associations were evaluated separately within treatment-naïve tumors using the full cohort available for each molecular layer: 485 tumors with matched RNA-sequencing and CNV data and 146 tumors with matched RNA-sequencing and DNA methylation data. Within each program, only program-member genes in the corresponding molecular matrix were tested. Program scores were modeled as a function of tissue site and the standardized gene-level purified logR or methylation M-value, with a patient-specific random intercept. Each gene was evaluated by likelihood-ratio comparison with a baseline model containing tissue site as a fixed effect and patient as a random intercept. *P* values were adjusted across tested genes within each program using the Benjamini–Hochberg procedure, and the standardized regression coefficient was used to represent the direction and magnitude of association.

### Assessment of CNV and DNA methylation contributions to anatomical variation

To determine whether anatomical differences in program activity were attenuated after accounting for intrinsic molecular variation, the same baseline, CNV-only, methylation-only and joint models were additionally fitted by restricted maximum likelihood. For each model, equal-weight estimated marginal means were calculated for tissue site; for models containing molecular covariates, CNV and/or methylation PCs were fixed at their sample means. Anatomical variance was defined as the variance across these site-specific estimated marginal means. Letting *V*_0_, *V_C_*, *V_M_*, and *V_CM_* denote the anatomical variance from the baseline, CNV-adjusted, methylation-adjusted and jointly adjusted models, respectively, the fractions of anatomical variance remaining were calculated as *V_C_* /*V*_0_, *V_M_*/*V*_0_ and *V_CM_*/*V*_0_. Accordingly, the fraction attenuated by joint molecular adjustment was 1 − V_CM_/V_0_. Values greater than one indicate that the adjusted anatomical-variance estimate exceeded the unadjusted estimate and were interpreted as showing no evidence of attenuation.

### Variance decomposition of transcriptional-program activity across tissue sites and patients

To quantify tissue-site and inter-patient heterogeneity in transcriptional-program activity, we analyzed treatment-naïve tumors and fitted a linear mixed-effects model for each of the 25 transcriptional programs. The model was specified as y_is_ = β_0_ + β_s_ + b_i_ + ε_is_, where y_is_ denotes the program score for a tumor from patient *i* and tissue site s, tissue site was included as a categorical fixed effect, and patient identity was included as a random intercept. Patient-specific intercepts and residual errors were assumed to follow independent normal distributions, b_i_ ~ N(0, σ^2^_patient_) and ε_is_ ~ N(0, σ^2^_residual_), respectively. Models were fitted by restricted maximum likelihood using lme4 ^44^.

To compare tissue-site and patient-associated heterogeneity on a common variance scale, tissue-site variance was calculated as the variance across the estimated marginal means for the four anatomical sites, with each site represented once irrespective of sample size: *V*_site_ = *Var*(μ^*_Adn_*, μ^*_Ome_*, μ^*_Per_*, μ^*_Asc_*). This quantity measures the dispersion of the modeled mean program activities across tissue sites under balanced site representation. Between-patient variance, *V*_patient_, was estimated as the patient random-intercept variance component, and residual variance, *V*_residual_, as the residual variance component of the fitted model. For each transcriptional program, the total variance under balanced site representation was defined as *V*_total,bal_ = *V*_site_ + *V*_patient_ + *V*_residual_. The tissue-site and patient variance proportions were then calculated as 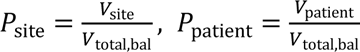.

Site-dependent transcriptional responses to chemotherapy For each transcriptional program, treatment-phase effects were assessed using a linear mixed-effects model with anatomical site, treatment phase and their interaction as fixed effects and patient as a random intercept: score ∼ tissue site × treatment phase + (1|patient). Models were fitted by maximum likelihood. For the reported pairwise comparisons, post-NACT versus treatment-naïve and relapse versus treatment-naïve, the model was refitted using only the two relevant phases and anatomical sites represented by at least three samples in each phase. Estimated marginal means, within-site phase differences and 95% confidence intervals were obtained using emmeans (version 1.11.1) ^45^. Effects were defined as the later-phase minus treatment-naïve difference. Within-site *P* values were adjusted across transcriptional programs within each anatomical site and phase comparison using the Benjamini–Hochberg method. For interaction plots, estimated marginal means were obtained from the full three-phase model.

Site dependence of the post-NACT response was assessed using interaction contrasts of within-site post-NACT versus treatment-naïve differences between each solid site (tubo-ovarian, omentum and peritoneum) and ascites.

### Cross-site inference of transcriptional compatibility and associated transcriptional programs

For each source–target pair, CSI was estimated using pairwise leave-one-patient-out linear discriminant analysis. Sample-level ICA component scores were first summarized by the median for each patient and anatomical site. For each held-out patient, the classifier was trained on tumors from the source and target sites from all other patients; centering, scaling and classifier fitting were performed using the training data only.

Within each training fold, all 21 batch-filtered ICA components were used as classifier features. Component scores were centered and scaled using means and standard deviations calculated from the training data, and the same transformation was applied to the held-out samples. Both site classes were required to contain at least two training observations. Linear discriminant analysis was fitted using equal source- and target-site class priors, preventing differences in site sample size from determining the posterior probabilities.

For a held-out source-site tumor, the posterior probability of assignment to the target-site class was interpreted as its source-to-target transcriptional compatibility. To reduce overconfident classification probabilities, posterior probabilities were temperature-scaled using (T=1.8): 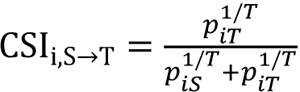, where *p_iS_* and *p_iT_* denote the untempered posterior probabilities for the source and target classes, respectively. Higher CSI values therefore indicate that the transcriptional state of the source-site tumor more closely resembles tumor states observed at the candidate target site in the remainder of the cohort.

To identify transcriptional programs associated with cross-site compatibility, we modeled CSI for each target site (omentum, peritoneum or ascites) as a function of program activity in tubo-ovarian tumors, fitting a separate univariable model for each program. For each program, we calculated the median score across each patient’s tubo-ovarian samples and z-standardized these patient-level scores across the patients included in the analysis. The primary association model was a bias-reduced beta regression with a logit link: *logit*(μ*_im_*) = α*_m_* + β*_m_Z_im_*, where μ_im_ denotes the expected transformed CSI value and *Z_im_* is the standardized activity of transcriptional program m in the tubo-ovarian tumor of patient *i*. CSI values were moved away from the boundaries of zero and one using the Smithson–Verkuilen transformation before model fitting. Heteroscedasticity-consistent HC2 covariance estimates were used for inference. When beta regression did not yield a valid estimate, ordinary least squares was fitted to logit-transformed CSI values using HC3 covariance estimates.

The reported effect size was the regression coefficient corresponding to a one-standard-deviation increase in program activity. Positive coefficients indicate greater compatibility with the candidate target site, whereas negative coefficients indicate lower compatibility. For each source–target comparison, nominal *P* values were adjusted across the tested transcriptional programs using the Benjamini–Hochberg method. Patient-level bootstrap resampling was performed in 2,000 replicates to assess the robustness of program associations and to estimate percentile confidence intervals for average marginal effects.

### Single-nucleus multiome sequencing

Nuclei were isolated from frozen samples using the Chromium Nuclei Isolation Kit (10x Genomics, #PN-1000494) according to the manufacturer’s instructions. The concentration and quality of the isolated nuclei were assessed using a LUNA-FX7™ Automated Cell Counter with Acridine Orange/Propidium Iodide stain (Logos Biosystems, #F23001). Paired chromatin-accessibility (ATAC) and gene-expression (GEX) libraries were generated using the Chromium Next GEM Single Cell Multiome ATAC + Gene Expression Reagent Bundle (10x Genomics, #PN-1000283), following the Chromium Single Cell Multiome ATAC + Gene Expression User Guide (CG000338). Briefly, nuclei were subjected to transposition and subsequently partitioned into Gel Beads-in-Emulsion (GEMs), enabling accessible chromatin fragments and nuclear RNA from the same nucleus to be indexed with a shared cellular barcode. The resulting ATAC and GEX products were processed separately to generate sequencing-ready libraries. The libraries were sequenced in paired-end mode (2 × 150 bp) on an Illumina NovaSeq X platform.

### Single-nucleus multiome data processing

Paired snRNA-seq and snATAC-seq libraries were generated using the 10x Genomics Chromium Single Cell Multiome platform and processed with Cell Ranger ARC v2.0.2 (10x Genomics). Aggregated feature-barcode matrices were imported into Python using Muon v0.1.7 ^46^ and separated into RNA and ATAC modalities for downstream analyses. All samples were initially processed jointly to generate a cohort-wide embedding, after which analyses were performed on the subset of samples included in this study.

For RNA quality control, nuclei with fewer than 100 detected genes, fewer than 500 or more than 15,000 UMIs, or greater than 20% mitochondrial transcripts were excluded. Doublets were identified independently within each sample using the R package scDblFinder v1.12.0 ^47^ and removed. RNA counts were normalized to 10,000 counts per nucleus, log-transformed, and 3,000 highly variable genes were identified using Scanpy v1.11.5. Principal component analysis, neighborhood graph construction, Leiden clustering, and UMAP visualization were performed using Scanpy workflows ^48^. Nuclei were annotated as cancer, immune or stromal cells using canonical markers and cell type signature scores derived from the reference atlas of Vázquez-García et al. ^12^.

ATAC profiles were processed using TF-IDF normalization followed by latent semantic indexing (LSI). The first LSI component, which predominantly captured sequencing depth, was excluded from downstream analyses. Because chromatin accessibility metrics varied substantially across major cellular compartments, ATAC quality-control thresholds were applied separately to RNA-defined cancer, immune, and stromal cells. Cancer and stromal cells were required to have >600 detected ATAC peaks and >1,000 ATAC fragments, whereas immune cells were required to have >200 detected peaks and >500 fragments. Cells passing ATAC quality control were subsequently filtered using compartment-specific RNA criteria.

Immune cells were required to have >300 detected genes and 500-15,000 UMIs, stromal cells >800 detected genes and 900-15,000 UMIs, and cancer cells >800 detected genes and 1,500-15,000 UMIs. All cells were required to have <15% mitochondrial transcripts.

Large-scale CNV were inferred from RNA expression profiles using infercnvpy v0.6.1 (https://github.com/icbi-lab/infercnvpy). Immune and stromal cells were used as diploid reference populations. Inferred copy-number profiles were used to distinguish malignant from non-malignant cells and we filtered out epithelial cells with inferred CNV profiles that clustered together with stromal cells.

### Classification of Mes-ECM and Mechano-EMP malignant-cell states

Malignant epithelial cells were stratified according to Mes-ECM and Mechano-EMP program activity. Per-cell program scores were calculated using UCell (version 2.2.0) ^49^. High-activity thresholds were determined independently for each program using knee-point detection on the ranked UCell-score distribution. Briefly, rank and score were normalized to [0,1], the perpendicular distance from each point to the line connecting the distribution endpoints was calculated, and the maximum distance after running-median smoothing (window = 5) and exclusion of the outer 5% of observations was used as the threshold. Cells were classified as dual-low, Mes-ECM-high, Mechano-EMP-high or dual-high according to the resulting binary program assignments.

### Differential chromatin accessibility and motif enrichment

Differential chromatin accessibility was assessed between Mes-ECM-high, Mechano-EMP-high or dual-high cells and dual-low cells using logistic regression implemented in FindMarkers in Seurat (version 5.3.0) ^50^, with total ATAC counts included as a latent variable. Gained peaks were defined by adjusted P < 0.01, average log2 fold change > 0.2 and detectable accessibility in >3% of cells in the corresponding program-high state; depleted peaks were defined analogously using an average log2 fold change < −0.2 and >3% accessibility in dual-low cells.

Motif enrichment was assessed separately among gained and depleted peaks using FindMotifs in Signac (version 1.14.0) ^51^. For comparative analyses, motif enrichment among gained and depleted peaks was represented by positive and negative signed log2 fold enrichment, respectively. Motifs with FDR < 0.05 and signed log2 fold enrichment > 0.2 in both the Mes-ECM-high and Mechano-EMP-high comparisons were classified as shared gain-associated motifs, whereas those with FDR < 0.05 and signed log2 fold enrichment < −0.2 in both comparisons were classified as shared depletion-associated motifs. Motifs meeting the positive criterion in only one comparison were classified as state-specific gain-associated motifs.

### TF activity and co-regulatory network analysis

Gene-regulatory networks and TF activities were inferred using LINGER (version 1.105) ^52^ from the matched single-nucleus RNA and ATAC data. RNA and regulatory-element profiles were aggregated by sample for LINGER training, mitochondrial genes were excluded, and the resulting matrices were processed against the LINGER reference network using the hg38 genome. Models were trained using a ReLU activation function, followed by inference of TF–regulatory-element, cis-regulatory and trans-regulatory relationships. Cell-level TF activities were calculated using regulon_sparse from the inferred cell-population regulatory network.

For differential TF-activity analysis, cell-level activities were averaged within sample–state groups to generate sample–state pseudobulks; groups containing fewer than 10 cells were excluded. Mes-ECM-high, Mechano-EMP-high and dual-high pseudobulks were separately matched to dual-low pseudobulks from the same sample. For each TF, matched activity differences were tested against zero using two-sided one-sample t-tests, equivalent to paired t-tests on the pseudobulk activities, with at least three matched pairs required. P values were adjusted across all TF–contrast tests using the Benjamini–Hochberg method. TFs with increased activity and global FDR < 0.05 in both the Mes-ECM-high versus dual-low and Mechano-EMP-high versus dual-low comparisons were used to construct the shared co-regulatory network.

For network construction, the LINGER trans-regulatory matrix was restricted to these shared activated TFs and positive TF–target regulatory weights. For each TF, approximately the strongest 20% of positive target weights were retained, with at least the 100 strongest targets retained where available. TF–TF co-regulatory similarity was calculated as the cosine similarity between the resulting target-regulatory profiles. The sparse network retained the five nearest neighbors of each TF together with TF pairs in the top 20% of global similarities and was visualized using a similarity-weighted Fruchterman–Reingold layout.

### Functional and pathway characterization of Mes-ECM and Mechano-EMP states

Activity of the YAP/TAZ mechanoregulation gene set WP_MECHANOREGULATION_AND_PATHOLOGY_OF_YAPTAZ_VIA_HIPPO_AND_NON HIPPO_MECHANISMS was quantified at single-cell resolution using UCell (version 2.2.0) ^49^. Scores were averaged within sample–state pseudobulks, requiring at least 10 cells per group, and each program-high state was compared with matched dual-low pseudobulks using two-sided paired t-tests.

To identify biological processes associated with chromatin-accessibility gains, genes positively linked to gained peaks were identified from Signac peak-to-gene links and intersected with genes upregulated in the corresponding program-high state. Upregulated genes were defined by adjusted P < 0.01, average log2 fold change > 0.2 and expression in >3% of cells; ribosomal protein genes were excluded. Over-representation analysis was performed using a one-sided hypergeometric test against MSigDB-derived gene sets restricted to 20–1,000 genes. P values were adjusted using the Benjamini–Hochberg method.

PROGENy pathway activities were inferred from normalized RNA expression using decoupleR (version 2.4.0) ^53^ and the human PROGENy network comprising the top 500 responsive genes per pathway. Cell-level activities were estimated using the multivariate linear-model method and averaged within sample–state pseudobulks containing at least 10 cells. Each program-high state was matched to the dual-low state from the same patient and tissue, and pathway-specific differences were tested using two-sided paired t-tests, requiring at least three matched pairs. Paired standardized effect sizes were calculated as d_z_ = Δ̅/s_Δ_, where Δ̅ and s_Δ_ denote the mean and standard deviation of the matched activity differences, respectively. P values were adjusted across all pathway–contrast tests using the Benjamini–Hochberg method.

### Cell–cell communication analysis

Cell–cell communication was inferred using CellChat (version 2.2.0) ^54^ with the human ligand–receptor database. Communication probabilities were estimated using the trimean method; interactions involving cell populations represented by fewer than 10 cells were excluded. Ligand–receptor interactions with CellChat P < 0.05 were considered significant. Cancer-to-TME and TME-to-cancer interactions were evaluated separately. For each direction and TME population, an interaction was considered differentially detected for Mes-ECM if the ligand–receptor pair was significant for both Mes-ECM-high and dual-high cancer cells but non-significant for the corresponding interaction involving dual-low cancer cells. Differentially detected Mechano-EMP interactions were defined analogously, requiring significance for both Mechano-EMP-high and dual-high cancer cells and non-significance for dual-low cancer cells. Interactions meeting both criteria were considered shared.

For each cancer state, TME population and signaling direction, differentially detected interactions were summarized by the number of ligand–receptor pairs and the summed CellChat communication probability. For interaction-type summaries, ligand–receptor pairs involving CAFs, endothelial cells, pericytes and macrophages were grouped by ligand–receptor identity into trophic/guidance, matrix adhesion/remodeling, mechanosensory SDC4, and vascular/inflammatory coupling categories.

### H&E WSI acquisition and management

Formalin-fixed paraffin-embedded (FFPE) tissue blocks were collected at initial diagnosis for both diagnostic and research purposes. The FFPE blocks were prepared by the Pathology Department of Turku University Hospital (diagnostic samples) or Histology core facility at the University of Turku’s Institute of Biomedicine, Finland (research samples). The H&E staining process for all blocks was conducted at the Pathology Department of Turku University Hospital. Auria Biobank (University of Turku) performed the image scanning using 20x magnification, and the WSIs were subsequently stored in the OMERO database.

### Panoptic segmentation of WSIs

We simultaneously segmented tissue regions and nuclei in WSIs using a Cellpose-based panoptic model implemented in Histolytics ^15,55^. The model used a multitask U-Net architecture with a shared encoder and separate decoder branches for nuclear and tissue-region segmentation ^56^. The encoder backbone was EfficientNetV2-L pretrained on ImageNet ^57^. The model was trained on semi-manually annotated HGSC H&E image crops acquired at 20× magnification, with dataset preparation and annotation described previously ^15^. Annotations distinguished cancer, stromal, immune and dead-cell nuclei, and six tissue-region classes: cancer, stroma, necrosis, omental fat, hemorrhage and serum. Training images were tiled into overlapping 224 × 224-pixel patches using a sliding window with a 32-pixel stride. Input patches were min–max normalized, and StrongAugment was applied during training ^58^. The model was trained for 40 epochs with a batch size of 10.

### Morphological feature extraction

#### Global and local stromal-cell composition

For whole-slide cell-type quantification, nuclei of each class were counted across the entire segmented slide, and the stromal-cell fraction was used as the global stromal-cell proportion. Local stromal composition was quantified in cumulative circular neighborhoods centered on cancer-cell centroids. For each cancer cell, the proportion of stromal cells among all neighboring cells was calculated within radii of 16, 32, 64, 128 and 256 μm, excluding the central cell from the denominator. Cancer cells without neighboring cells at a given radius were omitted. For each slide and radius, the local stromal-cell proportion was defined as the mean of these cancer-cell-centered proportions.

#### Quantification of collagen organization

Collagen features were quantified within segmented stromal regions. Nuclei were rasterized from cell-segmentation polygons and dilated by eight pixels to minimize detection of nuclear boundaries as collagen. Bright tissue-free regions were identified by grayscale thresholding at 220, followed by morphological closing, removal of objects smaller than 2,048 pixels and eight-pixel dilation. Pixels outside stromal regions or within nuclear masks or masks of bright tissue-free regions were excluded. Tiles were read with 50-pixel padding to reduce boundary artifacts, which was removed before feature extraction.

The remaining stromal image was converted to inverted grayscale, and collagen-associated edges were detected using a Canny edge detector with a Gaussian smoothing parameter of σ = 1.5 ^59^. Edges arising along boundaries between retained stromal regions and excluded areas were removed using a 5 × 5-pixel dilation–erosion boundary mask, and connected components smaller than 16 pixels were excluded. Each remaining component was treated as a collagen edge. Straight edges were defined by a minor-to-major axis-length ratio <0.2; collagen straightness was quantified as the fraction of detected edges classified as straight, and collagen-edge length by major-axis length.

Collagen orientation disorder was quantified among straight edges using a previously described orientation co-occurrence framework ^60,61^. Absolute orientations were assigned to 18 equally spaced bins spanning 0 to π/2, and a length-weighted orientation co-occurrence matrix was constructed, with contributions from each reference edge weighted by its major-axis length. After normalization to unit sum, collagen-fiber orientation-distribution entropy was calculated as H_CFOD_ = − ∑*_a_*_,*b*_ P*_ab_* log_2_ P*_ab_*, where P*_ab_* denotes an element of the normalized co-occurrence matrix.

#### Quantification of cancer-cell nuclear texture

Cancer-cell nuclear texture was quantified from hematoxylin optical-density images using nuclei identified by deep-learning-based panoptic segmentation. To account for inter-slide staining variation, slide-specific stain matrices were estimated using the Macenko method ^62^. Tissue-rich patches (>80% tissue) were sampled from tissue masks generated from slide thumbnails using Otsu thresholding ^63^. Patches intersecting annotated hemorrhage, serum or omental regions, containing <1% nuclear area, or containing >10% very dark or bright pixels (grayscale <50 or >220) were excluded; at least five valid patches were required per slide. RGB intensities were transformed to optical-density space, pixels with optical density >0.15 in all channels were retained, and two stain vectors were derived from the 10th and 90th percentiles of the angular distribution within the principal stain plane. The vectors were assigned to hematoxylin and eosin according to their relative enrichment within segmented nuclei.

To standardize stain intensity across slides, cohort-wide 1st and 99th percentiles of deconvolved hematoxylin and eosin intensities were estimated from 256-μm patches. For each slide, up to 50 patches containing ≥80% tissue and 1–80% nuclear area were sampled and deconvolved using the corresponding slide-specific stain matrix; patches with >80% very dark or bright pixels were excluded.

Nuclear texture was quantified in non-overlapping 256-μm image tiles containing ≥1% tissue. Hematoxylin intensities were obtained by optical-density color deconvolution using the slide-specific stain matrix, scaled to [0,1] using the cohort-wide 1st and 99th percentiles estimated from 256-μm patches, and Gaussian-smoothed σ = 0.5. Only nuclei fully contained within a tile and ≥20 pixels in area were analyzed. Within each nucleus, hematoxylin-dense patches were defined as connected pixels at or above the nucleus-specific 90th percentile of hematoxylin intensity; components smaller than five pixels were removed. For each nucleus, the number of retained hematoxylin-dense patches and the fraction of nuclear area occupied by the largest patch were quantified.

## DATA AVAILABILITY

Raw sequencing data have been deposited in the European Genome-phenome Archive (EGA) under the following study accession numbers: RNA sequencing (EGAS00001004714), whole-genome sequencing (EGAS00001006775) and whole-genome bisulfite sequencing (EGAS50000000727).

## ACKNOWLEDGEMENTS

This work was supported by the European Union’s Horizon 2020 Research and Innovation Programme under grant agreements 965193 (DECIDER) and 667403 (HERCULES); the Research Council of Finland (projects 363717, 325956 and 322927); the Sigrid Jusélius Foundation; Cancer Foundation Finland; the European Partnership for Personalised Medicine (EP PerMed) through the DYNAMITE project (Research Council of Finland funding decision 367829); and the Maud Kuistila Memorial Foundation. Digitized hematoxylin and eosin (H&E)-stained slides from archived samples collected for routine histopathological diagnostics were obtained from Auria Biobank. Figures 1A, 3N, 4E, 4F, 5H were created with Biorender.com.

## AUTHOR CONTRIBUTIONS

Conceptualization: K.Z., S.Ha., and A.V.; Methodology and Software: K.Z.; Formal Analysis: K.Z.; Validation: K.Z., E.K., and A.V.; Investigation: K.Z., L.F., D.A., E.K., G.M., Y.L., J.O., S.Hi., J.H., and A.V.; Resources: S.Hi., J.H., B.S., S.Ha., and A.V.; Data curation: K.Z., L.F., D.A., G.M., Y.L., J.O., J.H., B.S., S.Ha., and A.V.; Visualization: K.Z., D.A.; Supervision: S.Ha. and A.V.; Project Administration: S.Ha. and A.V.; Funding Acquisition: S.Hi., J.H., B.S., S.Ha., and A.V.; Writing - Original Draft: K.Z.; Writing - Review & Editing: all authors.

## DECLARATION OF INTERESTS

The authors declare no competing interests.

## Notes

### Competing Interest Statement

The authors have declared no competing interest.

### Summary of Updates

The manuscript has been substantially revised to include an expanded cohort of 600 specimens from 284 patients, with integrated analyses of RNA-seq, WGS and WGBS data. A reworked framework identifies recurrent cancer-cell transcriptional programs, and a new statistical framework links primary-tumor programs to solid and ascitic transcriptional phenotypes. Newly generated single-nucleus multiome data and expanded quantitative histomorphological analyses reveal shared and distinct regulatory and microenvironmental signaling patterns, together with associated features of tissue architecture. Analyses of genomic and epigenomic contributions, transcriptional remodeling following chemotherapy, and site-specific prognostic significance have also been updated. The title, text, figures and supplementary materials have been revised accordingly.

