## Supplementary Table S1 and Figures for "Transcriptional programs define solid and ascitic dissemination in ovarian cancer"

**Table S1. Description of cancer cell transcriptional programs.**

| Program Block | Transcriptional Program | Marker Genes | Representative Pathways | Description |
|---|---|---|---|---|
| **Cell cycle** | **Anabolic G1/S** | *CCNE2, E2F2, PHGDH, SLC7A5, SQLE* | mTORC1 signaling; E2F targets; G2/M checkpoint | Couples cell-cycle entry with anabolic and metabolic preparation. |
| | **Replication–Repair** | *RRM2, TICRR, TIMELESS, FANCI, BRIP1* | E2F targets; cell-cycle checkpoints; mitotic spindle checkpoint | Integrates DNA replication and repair with proliferative checkpoint activity. |
| | **CCNE1–FOXM1** | *CCNE1, FOXM1, CDK1, PLK1, AURKA* | E2F targets; G2/M checkpoint; mitotic spindle | CCNE1-linked proliferation coupled to FOXM1-driven mitotic progression. |
| | **Chromatin–Mitosis** | *HIST1H3B, HIST1H2BH, HIST1H1B, CDC20, BIRC5* | Mitotic cell cycle; prophase chromosome condensation; histone deacetylation | Links histone and chromatin regulation to mitotic execution. |
| **Interferon response** | **Canonical IFN** | *ISG15, MX1, OAS1, IFIT1, DDX58* | IFN-α response; IFN-γ response; interferon signaling | Canonical antiviral interferon-stimulated gene program. |
| | **IFN–Inflammatory** | *CXCL10, CXCL11, IRF1, NFKBIZ, TNFAIP3* | IFN-α response; IFN-γ response; TNF–NF-κB signaling | Interferon response coupled to inflammatory and chemokine signaling. |

| | | | | |
|---|---|---|---|---|
| | **IFN–MHC I** | *TAP1, TAP2, PSMB8, B2M, HLA-B* | Antigen cross-presentation; MHC-I peptide loading; MHC-I antigen processing | Interferon-associated antigen processing and MHC-I presentation. |
| **Mesenchymal states** | **Mesenchymal-ECM remodeling (Mes-ECM)** | *ZEB1, VIM, FN1, COL1A1, DDR2* | Epithelial–mesenchymal transition; ECM organization; collagen formation | Mesenchymal state characterized by extensive ECM deposition and remodeling. |
| | **Inflammatory mesenchymal (Inflam-Mes)** | *LOX, SERPINE1, MMP9, CCL2, GBP1* | Epithelial–mesenchymal transition; ECM organization; focal adhesion | Mesenchymal remodeling state with an inflammatory component. |
| | **Developmental mesenchymal (Dev-Mes)** | *SOX4, SOX11, HMGA2, CDH2, WNT7A* | Epithelial–mesenchymal transition; ECM organization; ECM proteoglycans | Developmentally patterned mesenchymal and ECM-remodeling state. |
| **Epithelial and cytoskeletal plasticity** | **Stress–Hypoxia** | *CA9, SLC2A1, ADM, STC2, TRIB3* | Hypoxia; keratinization; cornified-envelope formation | Stress-adaptive epithelial state associated with hypoxia and keratinization. |
| | **(Mechanoadaptive epithelial-mesenchymal plasticity) Mechano-EMP** | *SDC4, L1CAM, ITGB3, AMOTL2, FHL2* | Keratinization; cornified-envelope formation; early estrogen response | Mechanoadaptive epithelial–mesenchymal plasticity involving adhesion and polarity. |
| | **Epithelial Cohesion** | *TACSTD2, PERP, PPL, MPZL2, LAMA3* | Focal adhesion; extra-nuclear estrogen signaling | Epithelial structural state involving cohesion and cell–matrix adhesion. |

| | | | | |
|---|---|---|---|---|
| | **Rho–Cytoskeletal** | *ARHGEF17, ARHGEF5, ARHGAP23, CDC42BPG, MYH10* | RHO GTPase cycle; RHOA GTPase cycle; laminin interactions | Rho-regulated cytoskeletal, polarity, and architectural program. |
| **Inflammatory regulation** | **NF-κB–Inflammatory** | *NFKBIA, MYD88, S100A8, S100A9, CCL3* | TNF–NF-κB signaling; inflammatory response; interleukin signaling | Innate inflammatory program dominated by NF-κB and cytokine signaling. |
| | **Inflammatory Restraint** | *BCL6, ZBTB16, ZFP36, SGK1, PER1* | TNF–NF-κB signaling | Putative inflammation-modulating state involving regulatory restraint mechanisms. |
| **Lipid metabolism** | **Lipid Metabolism** | *PPARG, DGAT2, ACACB, LIPE, LPL* | Adipogenesis; hypoxia; TNF–NF-κB signaling | Lipid uptake, synthesis, storage, and mobilization program. |
| **Lineage and developmental states** | **Oncofetal Stemness** | *LIN28A, SALL4, TDGF1, SOX2, IGF2BP1* | Embryonic stem-cell pluripotency; pluripotency transcriptional regulation; gastrulation | Reactivation of embryonic, pluripotency-associated, and oncofetal programs. |
| | **Secretory Epithelium** | *HNF1A, HNF4A, PDX1, RBPJL, PTF1A* | Pancreatic acinar development; exocrine pancreas development; pancreatic progenitor lineage | Secretory epithelial differentiation with enteropancreatic features. |
| | **Motile Cilia** | *FOXJ1, RFX3, DNAI1, DNAH5, CCDC39* | Ciliopathies | Motile-cilia differentiation involving axonemal and dynein machinery. |

| | | | | |
|---|---|---|---|---|
| | **Skeletal Myogenesis** | *MYOD1, MYOG, ACTA1, MYH1, CKM* | Myogenesis; muscle contraction; striated-muscle contraction | Ectopic skeletal-muscle differentiation and contractile program. |
| | **HOX–Chromatin** | *HOXA10, HOXB13, HOXC13, HOXD13, HELLS* | Histone deacetylation; chromatin modification; DNA methylation | Developmental HOX-patterning program coupled to chromatin regulation. |
| **Translation and bioenergetics** | **Ribosomal Translation** | *RPL37, RPL35A, RPL38, RPL11, RPS24* | Translation elongation; translation initiation; cytoplasmic ribosomal proteins | Core ribosomal and cytoplasmic translation program. |
| | **Translation–OXPHOS** | *RPS13, RPL39, NDUFB5, COX5B, ATP5E* | Translation elongation; cotranslational protein targeting; cytoplasmic ribosomal proteins | Translation-associated program coupled to mitochondrial respiratory machinery. |
| | **Mitochondrial OXPHOS** | *PPARGC1B, MT-CO1, MT-CO2, MT-ND4, MT-ATP6* | Mitochondrial electron-transport chain; respiratory electron transport; aerobic respiration | Mitochondrial oxidative phosphorylation and respiratory electron transport. |

#### Figure S1

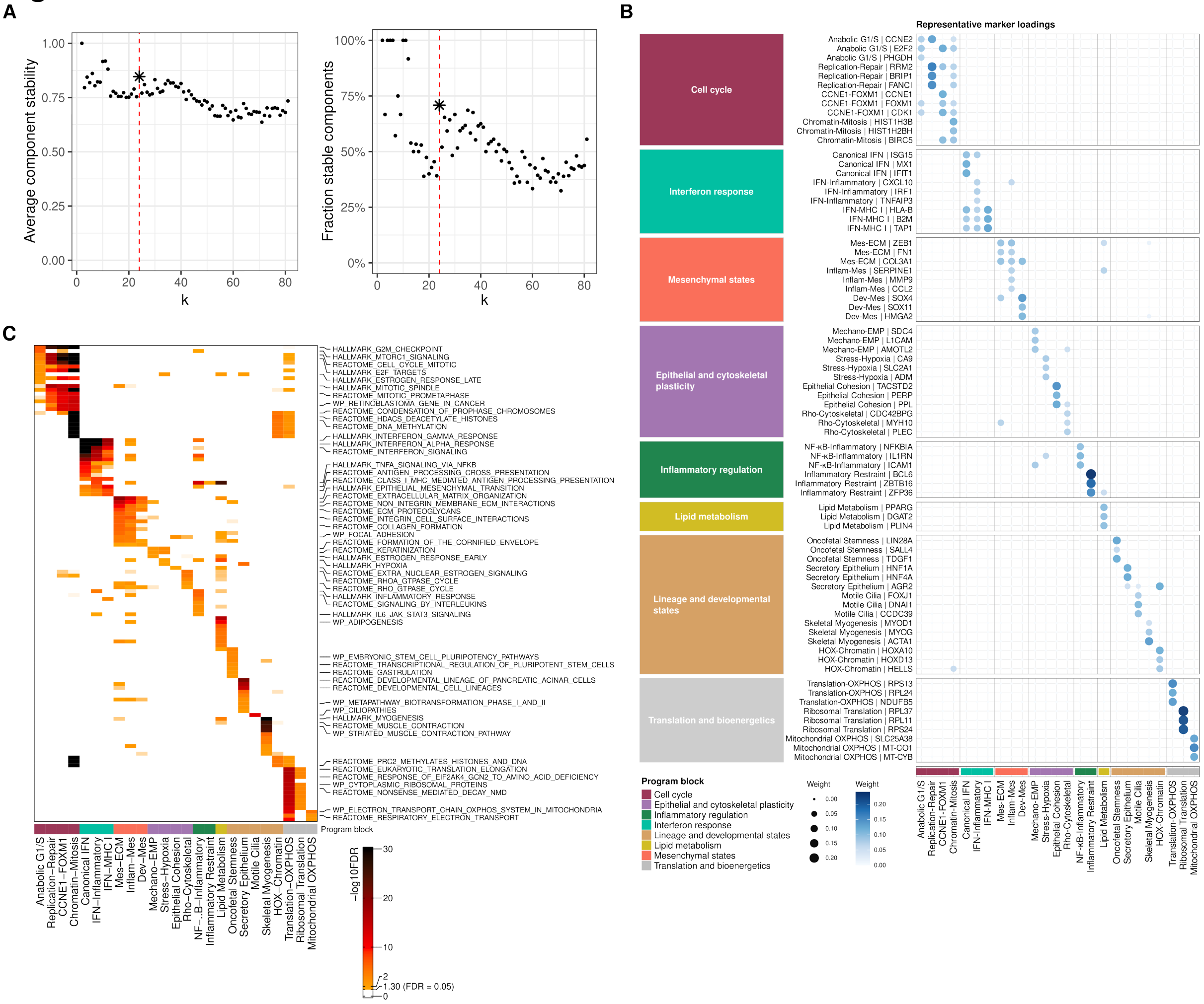




**Figure S1. Stability, marker specificity, and functional annotation of transcriptional programs. (A)** Stability of candidate ICA decompositions across the evaluated number of components, k. Left, average component stability; right, fraction of components with stability ≥0.75. The selected solution at k=24 is indicated by the red dashed line and asterisk. **(B)** Program loadings of representative marker genes for the 25 transcriptional programs. Three representative markers are shown for each program, and their weights are displayed across all programs. Dot size and color indicate gene weight. Markers and programs are grouped according to the eight broader biological categories indicated by the annotation blocks. **(C)** Gene-set enrichment profiles supporting functional annotation of the transcriptional programs. Color denotes enrichment significance as −log10(FDR). The annotation bar denotes the broader biological group assigned to each program.

#### Figure S2

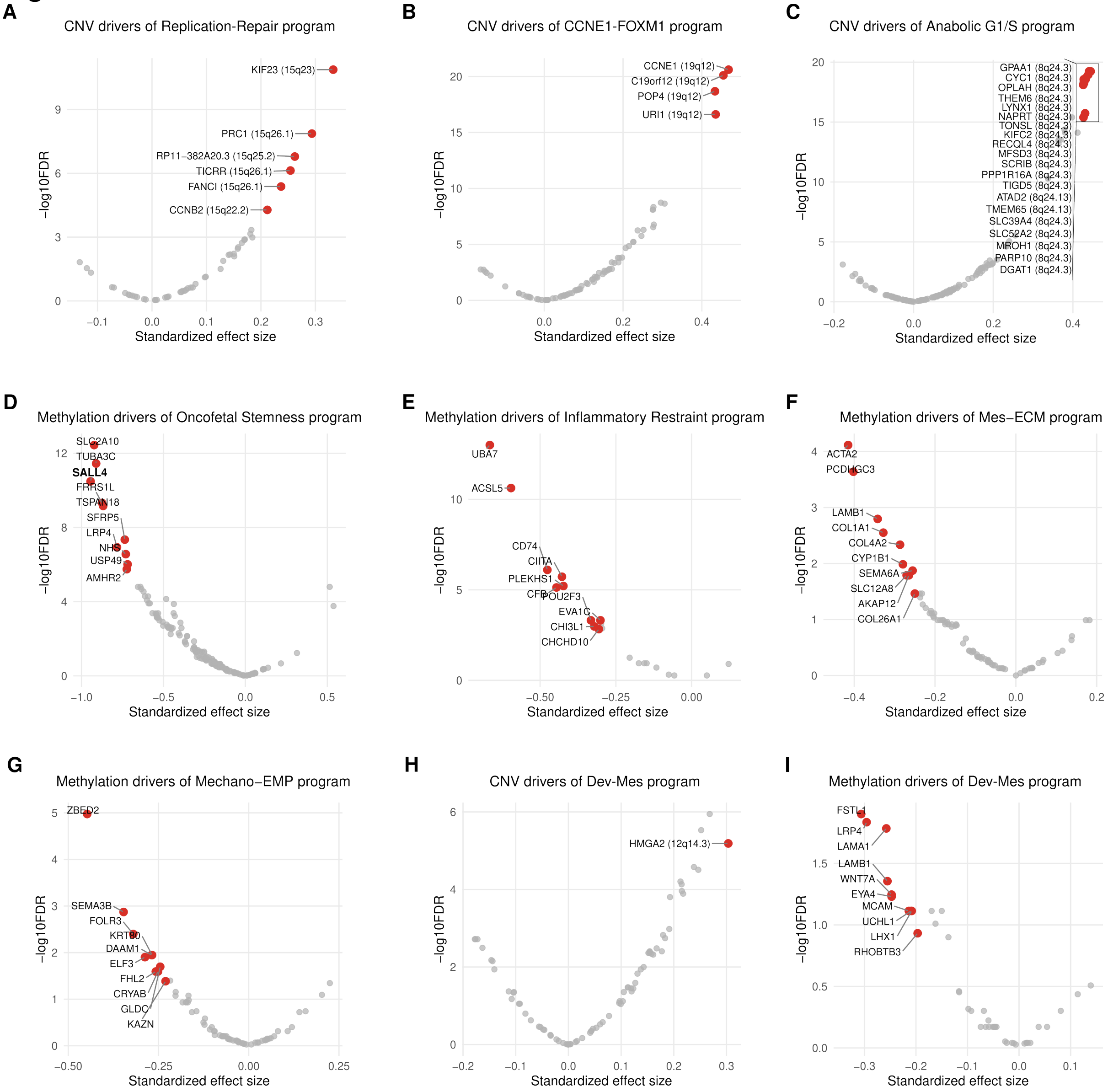


**Figure S2. Genetic and epigenetic determinants of transcriptional-program activity. (A–C)** Gene-level CNV associations for Replication–Repair (**A**), CCNE1–FOXM1 (**B**) and Anabolic G1/S (**C**). **(D–G)** Gene-level DNA-methylation associations for Oncofetal Stemness (**D**), Inflammatory Restraint (**E**), Mes-ECM (**F**), and Mechano-EMP (**G**) programs. **(H–I)** Gene-level CNV (**H**) and DNA-methylation (**I**) associations for Dev-Mes. CNV and methylation analyses included 485 and 146 treatment-naïve tumors, respectively, with matched expression data. Each point represents a gene; axes show standardized effect sizes and $-\log_{10}$(FDR) from linear mixed-effects models accounting for tissue site and patient identity. Selected genes are highlighted in red and labeled, with cytogenetic locations included in CNV panels.

### Figure S3

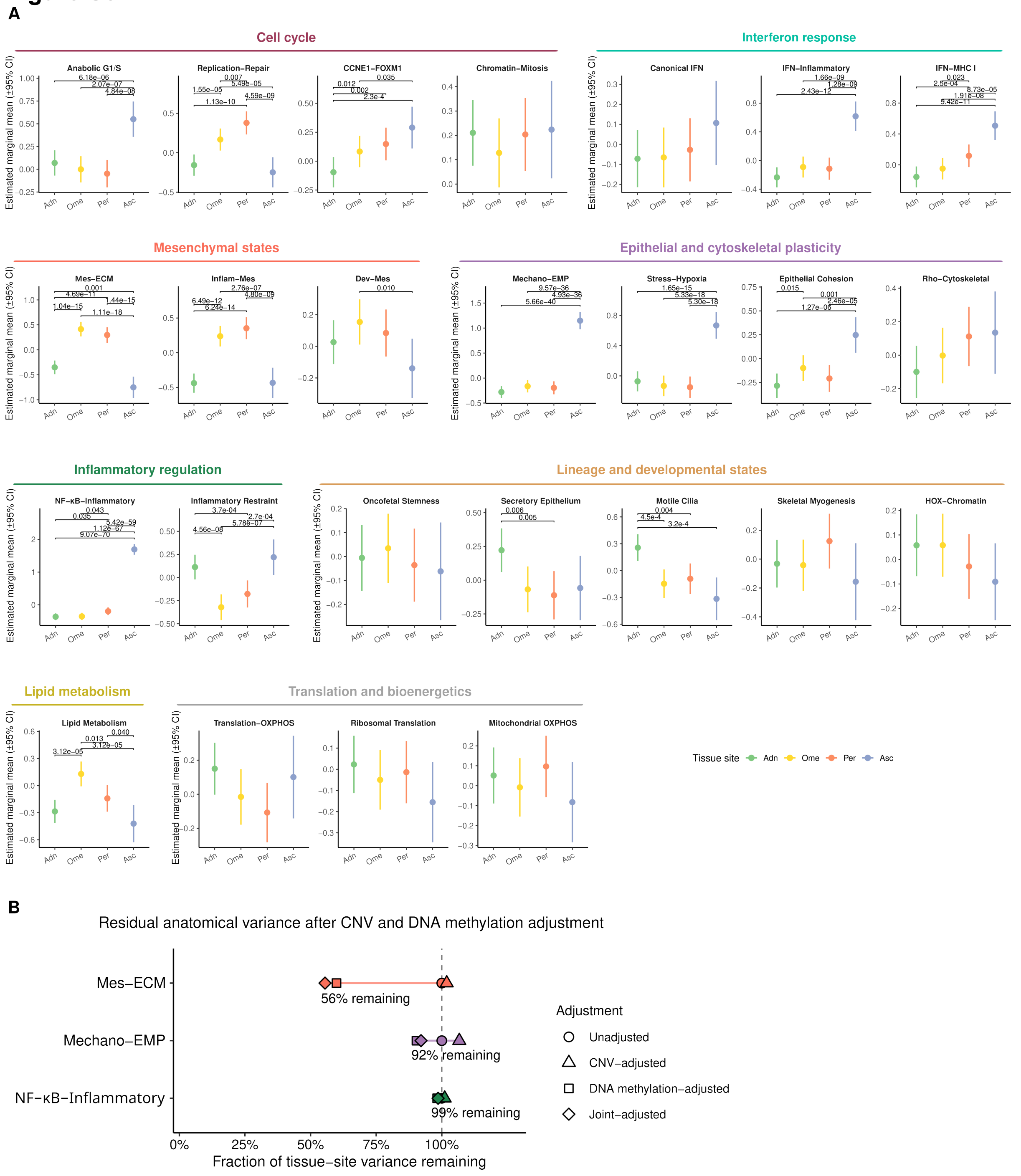


**Figure S3. Anatomical variation in transcriptional-program activity in treatment-naïve tumors. (A)** Estimated marginal mean (EMM) activity of each transcriptional program across tubo-ovarian tumors (Adn), omental metastases (Ome), peritoneal metastases (Per), and malignant ascites (Asc). Points indicate EMMs and error bars indicate 95% confidence intervals. Programs are grouped by program class. Pairwise differences between anatomical sites with BH-adjusted $P < 0.05$ are denoted. **(B)** Residual tissue-site variance for Mes-ECM, Mechano-EMP and NF-κB-Inflammatory after adjustment for CNV, DNA methylation or both, normalized to the unadjusted tissue-site variance for each program. Values greater than 1 indicate that the adjusted estimate exceeded the unadjusted estimate and were interpreted as no evidence of attenuation.

#### Figure S4

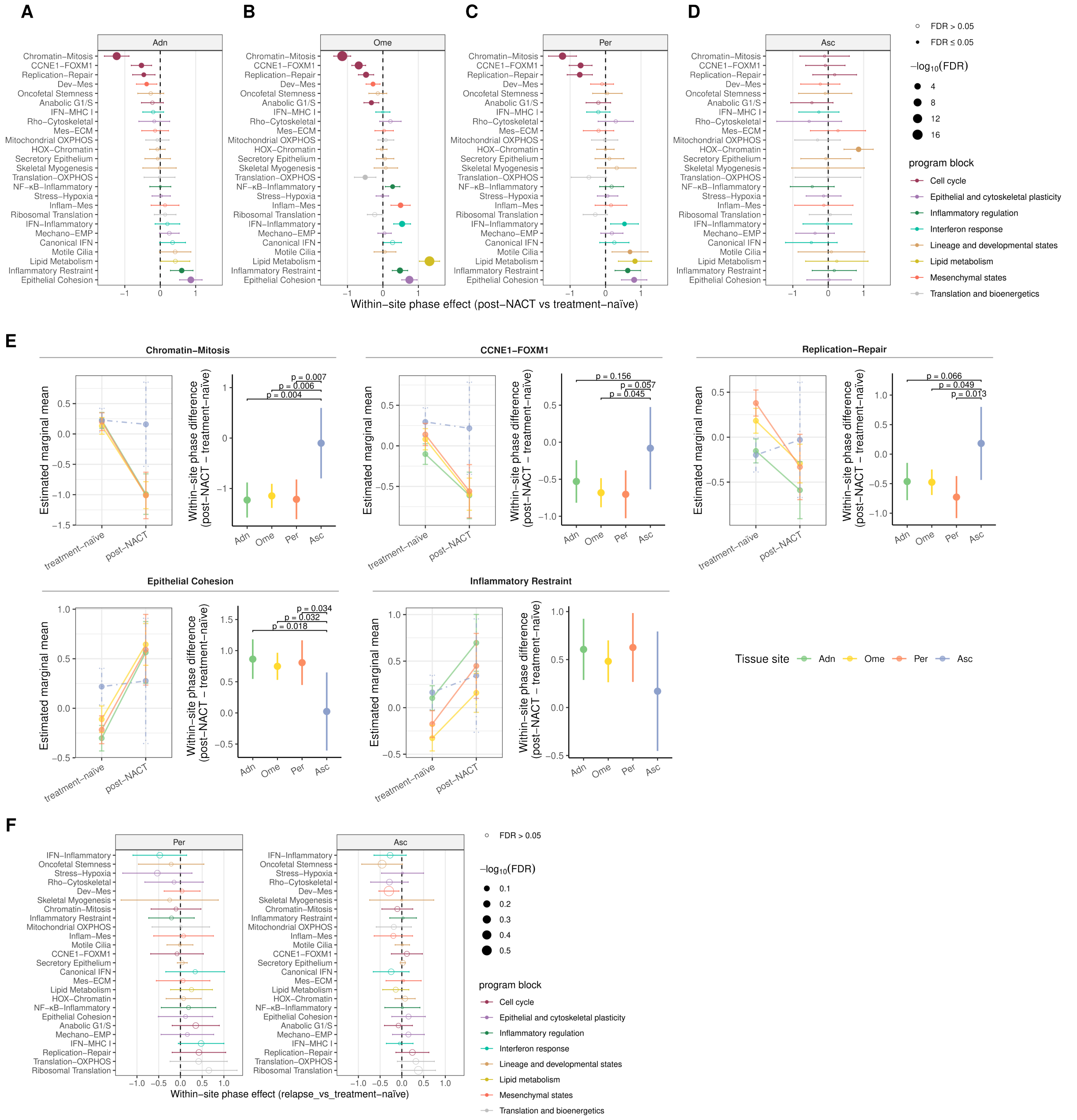


**Figure S4. Anatomical site shapes chemotherapy-associated transcriptional remodeling**. **(A–D)** Within-site post-NACT versus treatment-naïve differences in transcriptional-program activity in tubo-ovarian tumors (Adn; **A**), omental metastases (Ome; **B**), peritoneal metastases (Per; **C**), and malignant ascites (Asc; **D**). Points show the estimated differences, with positive values indicating higher activity in post-NACT tumors and horizontal bars indicating 95% confidence intervals. Point size represents −log10(FDR); filled points indicate FDR ≤ 0.05 and open points indicate FDR > 0.05. Colors denote transcriptional-program class. **(E)** Site-dependent post-NACT responses for five transcriptional programs showing significant post-NACT-associated changes in all three solid tumor sites but not in ascites. For each program, the left plot shows estimated marginal mean activity in treatment-naïve and post-NACT tumors across anatomical sites, and the right plot shows the corresponding within-site post-NACT versus treatment-naïve differences, with error bars indicating 95% confidence intervals. P values < 0.2 are shown for difference-of-differences contrasts comparing the post-NACT-associated change at each solid tumor site with that in ascites. **(F)** Within-site relapse versus treatment-naïve differences in transcriptional-program activity in peritoneal and ascitic tumors. Points show the estimated differences, with positive values indicating higher activity in relapse tumors and horizontal bars indicating 95% confidence intervals. Point size represents −log10(FDR); filled points indicate FDR ≤ 0.05 and open points indicate FDR > 0.05. Colors denote transcriptional-program class.

#### Figure S5

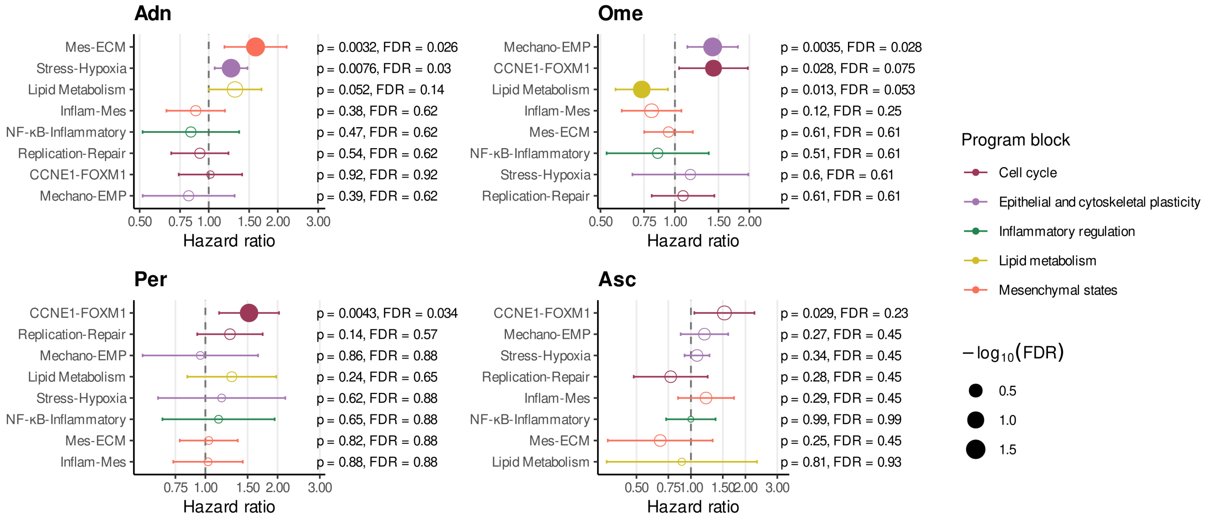


**Figure S5. Covariate-adjusted associations between dissemination-associated transcriptional programs and overall survival across anatomical sites.** Site-specific multivariable Cox models assessed associations between pretreatment activity of dissemination-associated transcriptional programs and overall survival for tubo-ovarian tumors (Adn), omental metastases (Ome), peritoneal metastases (Per), and malignant ascites (Asc). Models were adjusted for treatment strategy, age, FIGO stage, residual disease after cytoreductive surgery, and SBS3 status, used as a proxy for homologous recombination deficiency. Points indicate hazard ratios and horizontal lines indicate 95% confidence intervals; hazard ratios greater than 1 indicate association with shorter overall survival. Point colors denote program class and point sizes represent −log10(FDR). Filled points indicate associations with nominal P≤0.05 and FDR ≤0.20; nominal P values and FDRs are shown alongside each program.

#### Figure S6

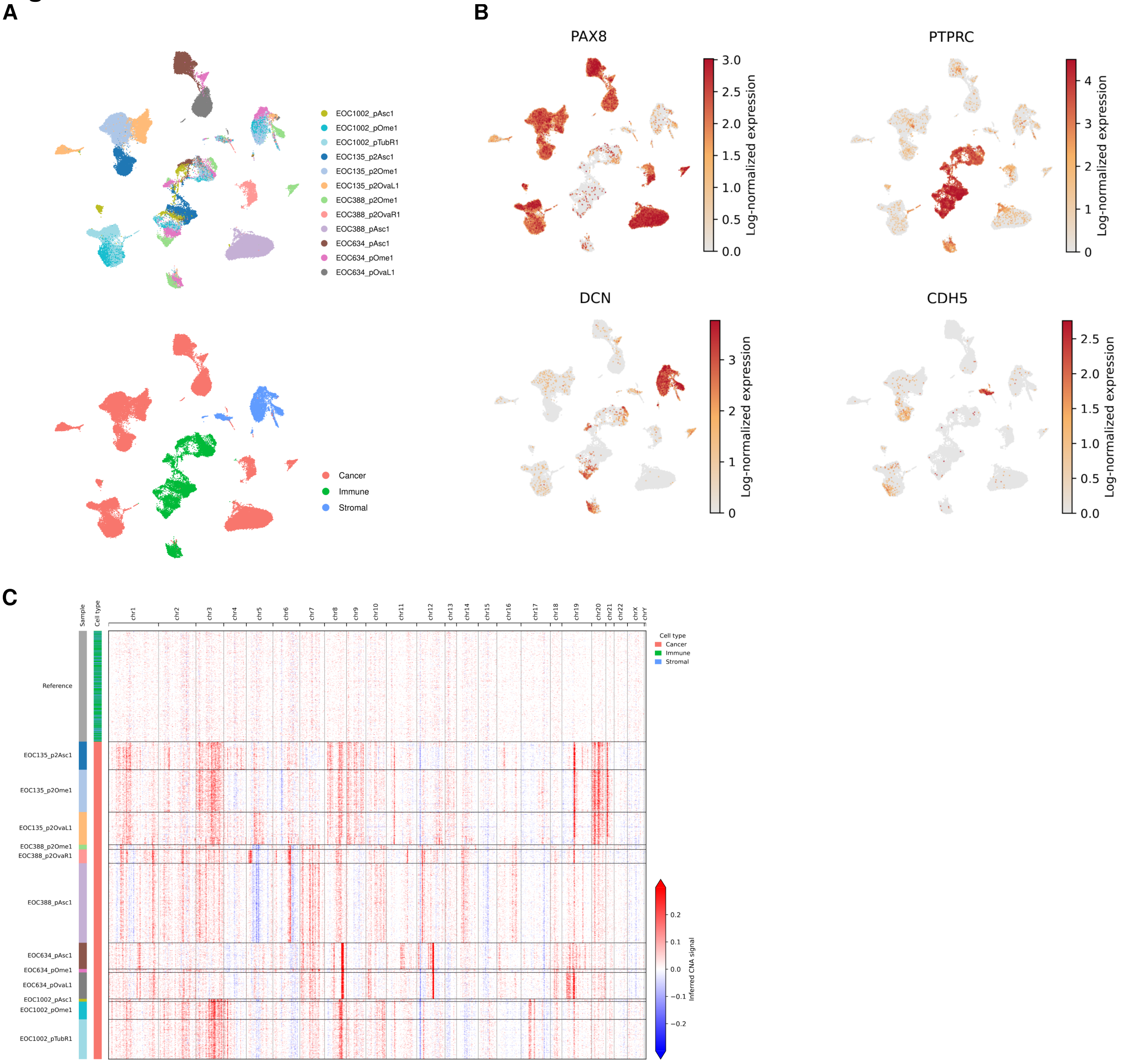


**Figure S6. Single-nucleus multiome landscape and major cell-type annotation. (A)** UMAP representation of nuclei profiled by single-nucleus multiome sequencing, colored by sample (top) and cell type (bottom). **(B)** UMAP plots showing the expression of canonical markers of malignant epithelial, stromal, and immune cells. **(C)** Inferred copy-number alteration (CNA) profiles.

#### Figure S7

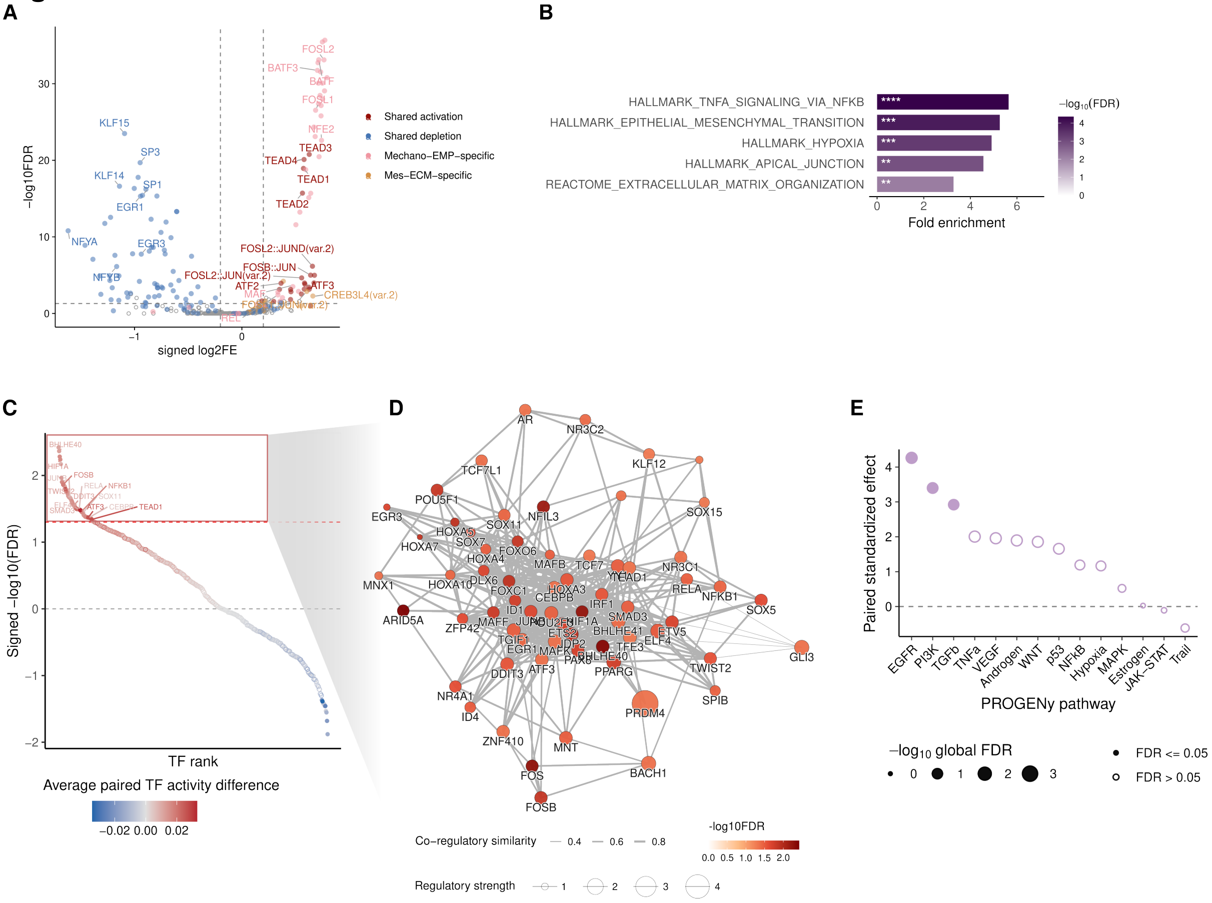


**Figure S7. Dual-high cells integrate Mes-ECM and Mechano-EMP regulatory programs. (A)** Motif enrichment among differentially accessible peaks in dual-high versus dual-low cells. Each point represents a motif. Colored points indicate motifs classified from the Mes-ECM-high versus dual-low and Mechano-EMP-high versus dual-low comparisons as shared activation, shared depletion, Mechano-EMP-specific, or Mes-ECM-specific; other motifs are shown as open grey points. Dashed lines indicate FDR = 0.05 and |log2 FE| = 0.2. **(B)** Representative gene sets enriched among upregulated genes positively linked to peaks with increased accessibility in dual-high versus dual-low cells. Bar length indicates fold enrichment, and fill indicates −log10(FDR). Asterisks indicate FDR thresholds: ∗FDR < 0.05, ∗∗FDR < 0.01, ∗∗∗FDR < 0.001, and ∗∗∗∗FDR < 0.0001. **(C)** Rank plot of differential TF activity in dual-high versus dual-low cells. TFs are ordered by signed −log10(FDR), with positive and negative values indicating increased and decreased activity in dual-high cells, respectively. Point color indicates the mean paired TF-activity difference, and filled and open points denote FDR < 0.05 and > 0.05, respectively. Selected regulators are labeled. **(D)** TF–TF co-regulatory similarity network of TFs with significantly increased activity in dual-high cells relative to dual-low cells (FDR < 0.05). Node fill indicates −log10(FDR), node size indicates regulatory strength, and edge width indicates co-regulatory similarity. **(E)** Differential PROGENy pathway activity in dual-high versus dual-low cells. Each point represents a pathway; the y-axis shows the paired standardized effect size calculated from the matched sample–state pseudobulk activity differences. Point size indicates −log10(FDR); filled and open points denote FDR ≤ 0.05 and > 0.05, respectively. Statistical analysis is as in Figure 3K.

### Figure S8

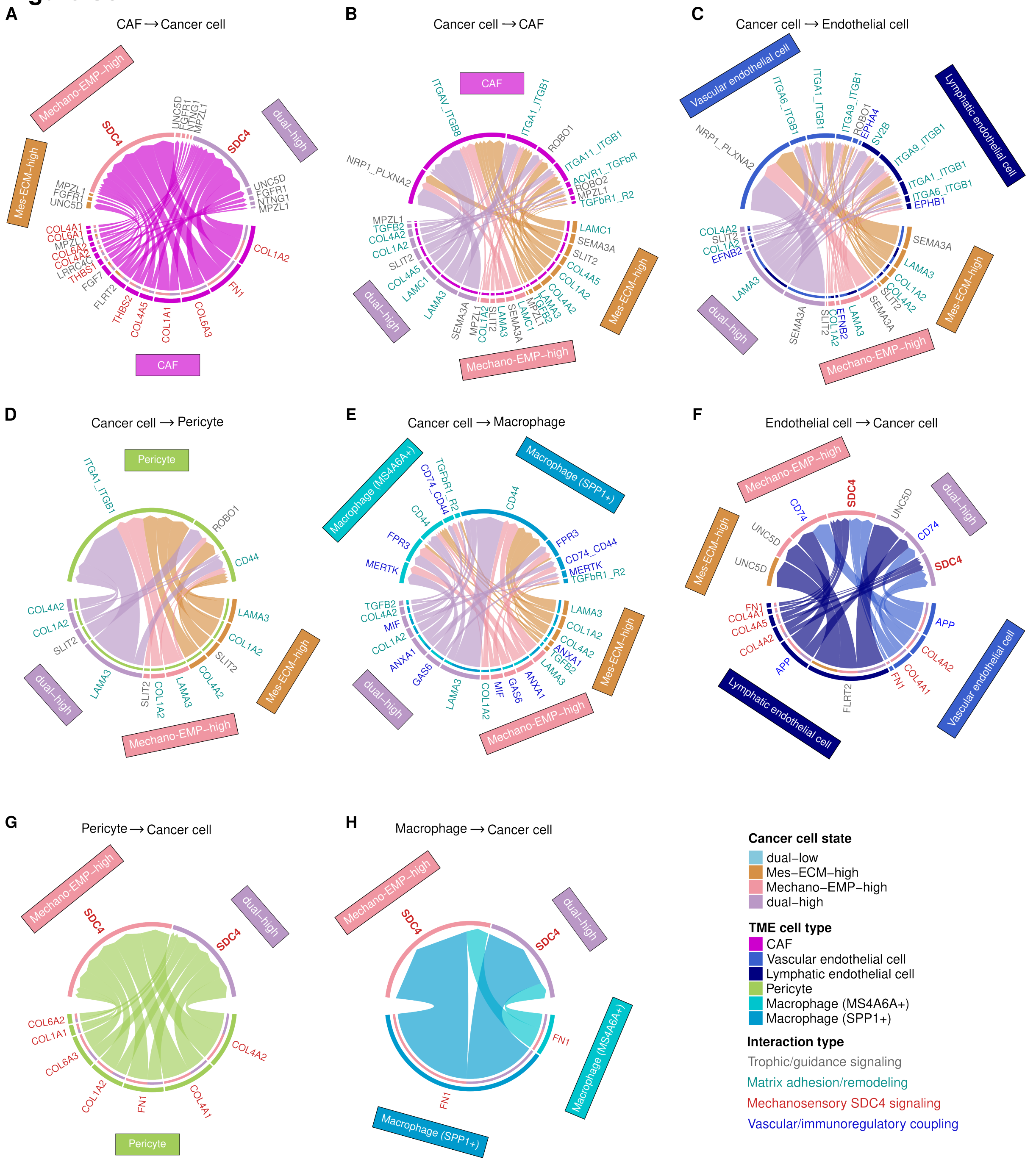


**Figure S8. Bidirectional tumor–microenvironment interactions associated with Mes-ECM and Mechano-EMP cancer-cell states.** Chord diagrams show ligand–receptor interactions from CAFs to cancer cells (**A**); from cancer cells to CAFs (**B**), vascular and lymphatic endothelial cells (**C**), pericytes (**D**) and macrophages (**E**); and from vascular and lymphatic endothelial cells (**F**), pericytes (**G**) and macrophages (**H**) to cancer cells. Interactions were retained if significant in at least one single-high state (Mes-ECM-high or Mechano-EMP-high) and also in dual-high cells ($P < 0.05$), but non-significant for the corresponding interaction involving dual-low cells. Dual-low cells are therefore not shown. Sector colors denote cancer-cell states or TME populations, chord colors denote sender populations, and gene-label colors indicate interaction categories. CAFs, cancer-associated fibroblasts.

#### Figure S9

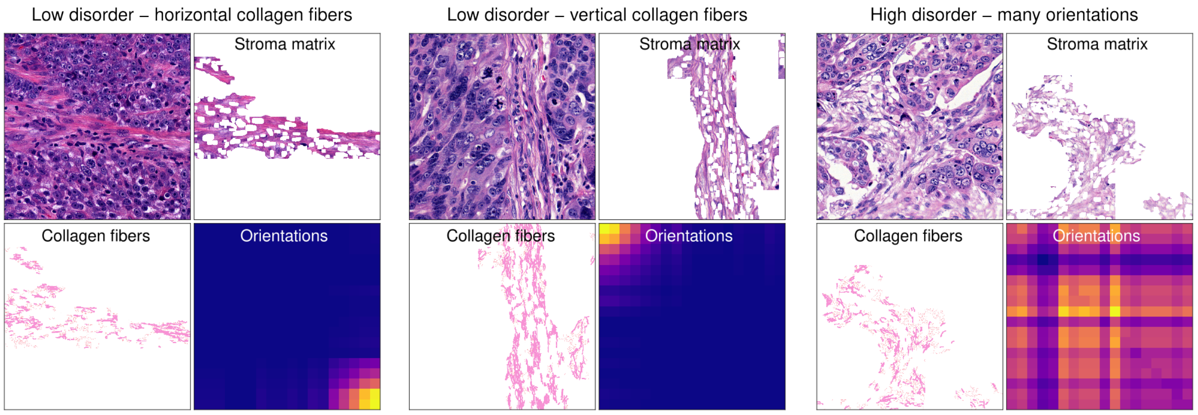


**Figure S9. Representative patterns of collagen-fiber organization.** Examples illustrating stromal matrix regions, detected collagen fibers, and corresponding orientation co-occurrence matrices for regions with predominantly horizontal fibers (left), predominantly vertical fibers (middle), or fibers distributed across multiple orientations (right).

#### Figure S10

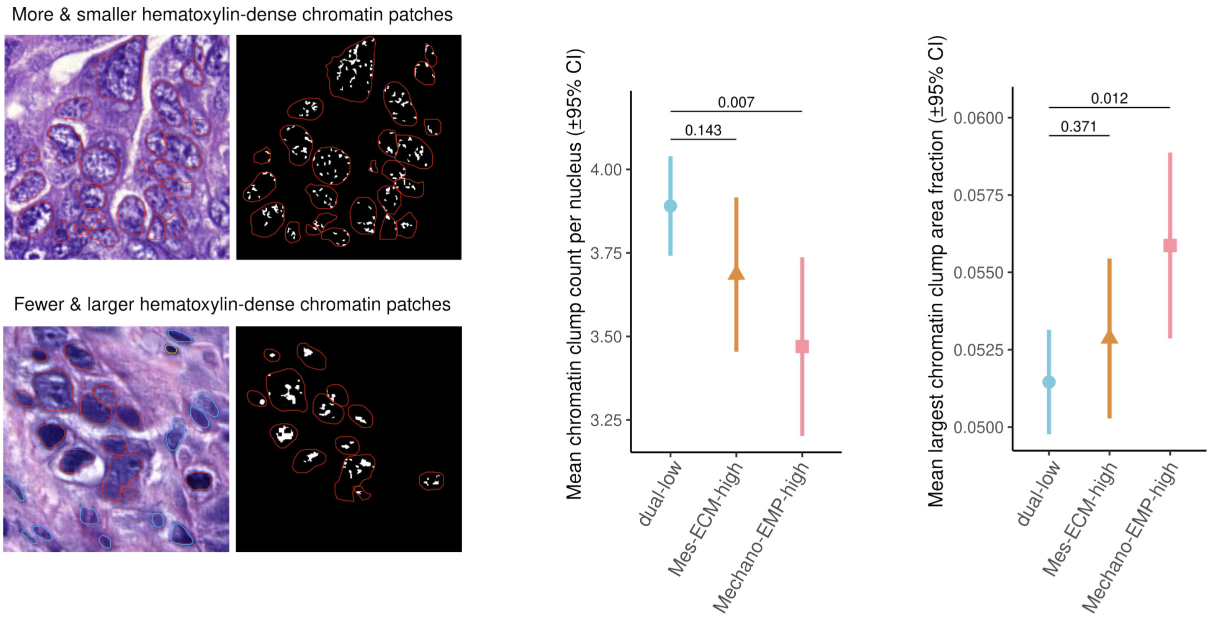


**Figure S10. Cancer-cell nuclear texture across transcriptional states.** Left, representative H&E images and corresponding detected hematoxylin-dense intranuclear patches illustrating nuclei with more and smaller or fewer and larger dense patches. Middle, EMM number of hematoxylin-dense patches per cancer-cell nucleus with 95% CIs. Right, EMM fraction of nuclear area occupied by the largest hematoxylin-dense patch with 95% CIs. P values for comparisons with dual-low tumors are indicated.
